# SRSF10 is essential for progenitor spermatogonia expansion by regulating alternative splicing

**DOI:** 10.1101/2022.03.06.483179

**Authors:** Wenbo Liu, Xukun Lu, Zheng-Hui Zhao, Qian-Nan Li, Yue Xue, Zheng Gao, Si-Min Sun, Wen-Long Lei, Lei Li, Geng An, Hanyan Liu, Zhiming Han, Ying-Chun Ouyang, Yi Hou, Zhen-Bo Wang, Qing-Yuan Sun, Jianqiao Liu

## Abstract

Alternative splicing expands the transcriptome and proteome complexity and plays essential roles in tissue development and human diseases. However, how alternative splicing regulates spermatogenesis remains largely unknown. Here, using germ cell-specific knockout mouse model, we demonstrated that the splicing factor *Srsf10* is essential for spermatogenesis and male fertility. Depletion of *Srsf10* in germ cells had little effect on the formation of SSCs but impeded the expansion of progenitor spermatogonia, leading to the failure of spermatogonia differentiation and meiosis initiation. This was further evidenced by the decreased expression of progenitor cell markers in bulk RNA-seq, and much less progenitor and differentiating spermatogonia in single-cell RNA-seq data. Furthermore, the expression of genes involved in cell cycle was abnormal in all subtypes of spermatogonia identified in single-cell RNA-seq data. Notably, using isolated spermatogonia, we found that *Srsf10* depletion disturbed the alternative splicing of hundreds of genes, which were preferentially associated with cell cycle, mitotic cell cycle checkpoint and germ cell development, including *Dazl*, *Kit*, *Ret*, *Sycp1*, *Nasp* and *Bor*a. These data suggest that SRSF10 is critical for the expansion of progenitor spermatogonia by regulating alternative splicing, expanding our understanding of the mechanism underlying spermatogenesis.

## Introduction

Spermatogenesis is a complex and highly coordinated process during which spermatogonial stem cells (SSCs) give rise to haploid spermatozoa sustainably throughout life. The balance of self-renewal and differentiation of SSCs is fundamental for maintenance of spermatogenesis throughout life. Over self-renewal of SSCs will lead to stem cell accumulation and impair spermatogenesis, and even induce tumor. Conversely, over-differentiation will lead to SSC exhaustion, and thus progressive loss of germ cells and Sertoli cell-only syndrome^1^. Around embryonic days 15 (E15) in mice, male germ cells do not enter meiosis but are arrested at the G0/G1 phase, and are referred to as pre-spermatogonia. The pre-spermatogonia resume mitotic proliferation after birth and migrate from the center to the peripheral basement membrane, entering the appropriate environment (stem cell niche) to develop into SSCs. SSCs can self-renew to sustain the stem cell pool or differentiate to generate progenitor cells destined to differentiation^2, 3^. The stem cells and progenitor spermatogonia are collectively called undifferentiated spermatogonia which include A-single (As, single cells), A-paired (Apr, 2 cells interconnected by cytoplasmic bridges) and A-aligned (Aal, 4, 8 or 16 cells interconnected by cytoplasmic bridges) spermatogonia. Then, Aal spermatogonia transform into type A1 spermatogonia and further differentiate into A2, A3, A4, intermediate (In) and B spermatogonia. All these cell types are identified as differentiating spermatogonia^4^. Then, type B spermatogonia will divide into pre-leptotene spermatocytes which are the last stage of the mitotic phase and have the competence to enter meiosis. After two rounds of meiotic divisions, haploid spermatids are formed^5^. A large number of genes and multiple regulatory layers of gene expression, including transcriptional and post-transcriptional regulation are reported to be involved in such a complex process to ensure successful spermatogenesis. However, much remains to be understood regarding the homeostasis of SSCs in this intricate process.

Alternative splicing (AS) is a very important and universal post-transcriptional regulatory mechanism to expand the diversity of transcripts and proteins from a limited number of genes^6^. Importantly, AS occurs more frequently in higher mammals (90-95% of human genes) and complex organs (brain, heart and testes)^7, 8^, suggesting that AS contributes to the complexity of the organism. Large scale analysis based on Expressed Sequence Tags (ESTs) and deep sequencing revealed that AS was at an unusually high level in testes, where the transcription of the genome is substantially more widespread than in other organs^9, 10, 11^, indicating the regulatory functions of AS in the development of testes. Recently, single-cell RNA sequencing (scRNA-seq) revealed that the Gene Ontology (GO) terms of mRNA splicing and mRNA processing were enriched in the spermatogonia^12^. Moreover, many RNA splicing proteins were more highly expressed in type A spermatogonia and in pachytene spermatocyte cell clusters^13, 14^. AS isoform regulation impact greatly on the germ cell transcriptome as cells transit from the mitotic to meiotic stages of spermatogenesis^15^, suggesting that AS may be very effective for the mitotic division of spermatogonia. Recently, the RNA helicase DDX5 is shown to play essential post-transcriptional roles in the maintenance and function of spermatogonia via regulating the splicing of functional genes in spermatogonia^16^. *Mettl3*-mediated m6A regulates the alternative splicing of genes functioning in spermatogenesis and is critical for spermatogonial differentiation and meiosis initiation^17^. Our previous study has also revealed that *Bcas2* regulates AS in spermatogonia and is involved in the meiosis initiation^18^. Despite these encouraging findings, our knowledge of the mechanistic regulation of spermatogenesis, such as what AS factors are involved in this complicated process is still very limited.

AS is regulated by cis-regulatory sequences in pre-mRNAs and trans-acting splicing factors which are mostly RNA-binding proteins that bind to the cis-regulatory sequences and regulate splice site selection^19, 20^. Serine and arginine-rich (SR) proteins are one of the classic trans-acting splicing regulators which contain one or two RNA-recognition motifs (RRM) in the N-terminus and arginine/serine amino acid sequences (RS domain) in the C-terminus^21^. SR proteins are often identified as positive splicing regulators that promote exon inclusion through interacting with many other splicing factors^21, 22^. SRSF10 is an atypical SR protein functioning as a general splicing repressor when dephosphorylated^23^. In vitro assays show that SRSF10 is dephosphorylated to inhibit the splicing in mitotic cells or response to heat shock^24, 25^. In vivo, *Srsf10* knockout mice survived only until E15.5 due to multiple cardiac defects as a result of dysregulation of cardiac-specific alternative splicing of triadin pre-mRNA which is required for Ca^2+^ handling in the embryonic heart^26^. SRSF10 also plays important roles in myoblast differentiation, glucose production and adipocyte differentiation by regulating the correct functional alternative splicing^27, 28, 29^. Thus, *Srsf10* is critical for many physiological processes, but the role of *Srsf10* in male germ cell development has not been elucidated.

In this study, we generated germ cell conditional *Srsf10* knockout mice and found that *Srsf10* was essential for spermatogenesis and male fertility. Depletion of *Srsf10* in germ cells impeded the expansion of progenitor spermatogonia, leading to the failure of efficient differentiation of spermatogonia and meiosis initiation. Single-cell RNA-seq data confirmed that progenitors and differentiating spermatogonia were seriously lost in the *Srsf10* knockout testes at P8. The cell cycle, proliferation and survival were impaired in the residual *Srsf10* depleted undifferentiated spermatogonia. Further analysis showed that SRSF10 was involved in alternative splicing of genes functioning in the cell cycle and germ cell development in undifferentiated spermatogonia. Our data reveal that SRSF10 is involved in the alternative splicing of spermatogonia and male fertility.

## Results

### *Srsf10* is essential for male fertility and spermatogenesis

To explore the function of *Srsf10* in male fertility and spermatogenesis, we mated *Srsf10*^Floxed/Floxed^ (*Srsf10*^F/F^) mice with *Vasa-Cre* transgenic mice in which the *Cre* recombinase driven by a *Vasa* promoter is specifically expressed in germ cells as early as embryonic day 15 (E15)^30^ (Figure 1A). *Srsf10^F/-^;Vasa-Cre* (*Srsf10^cKO^*) mice were used as experiment mice, while *Srsf10^F/+^;Vasa-Cre* mice were normal and used as the control in the following experiments. SRSF10 and MVH, a germ cell-specific marker, were co-stained and the results showed that SRSF10 expression was barely detected in MVH-positive cells of postnatal day 8 (P8) *Srsf10*^cKO^ testes (Figure 1B), indicating that *Srsf10* was specifically depleted in germ cells as early as P8.

**Figure 1.**
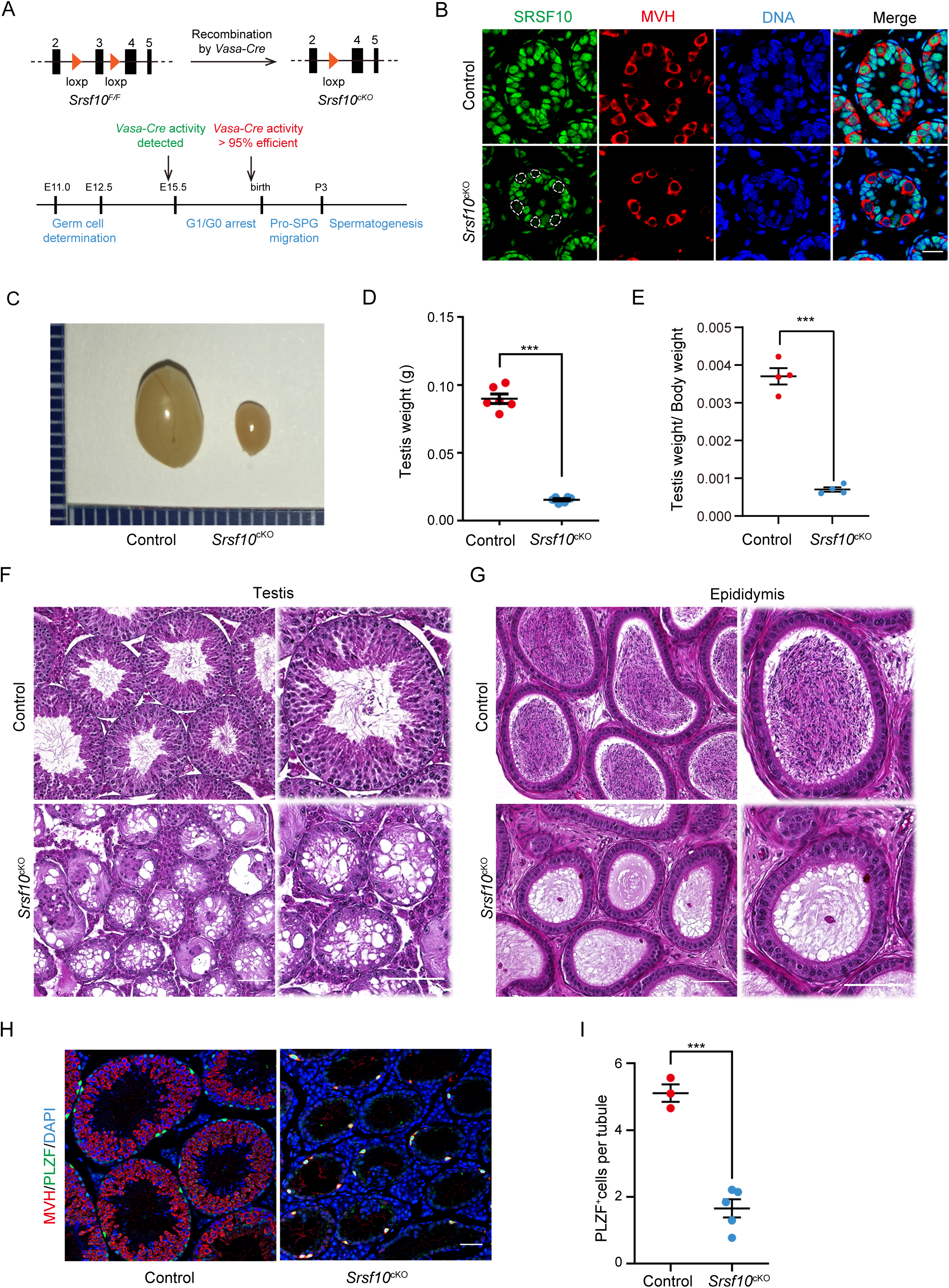
*Srsf10* is required for spermatogenesis and male fertility. A. Schematic diagram showing the deletion of *Srsf10* exon 3 and generation of *Srsf10*^cKO^ by *Vasa-Cre* mediated recombination in male germ cells as early as E15. B. IF staining for SRSF10 in the control and *Srsf10*^cKO^ testes of P8 mice. White circles denote the SRSF10 null germ cells. MVH (a germ cell marker) was co-stained to indicate the location of germ cells. The DNA was stained with Hoechst 33342. Scale bar, 20 μm. C. Morphological analysis of the adult testes of control and *Srsf10*^cKO^ mice. D. Testis weight of adult control and *Srsf10*^cKO^ mice (\*\*\**P* < 0.001, n = 6). Error bars represent s.e.m. E. The ratio of testes to body weight in adult control and *Srsf10*^cKO^ mice (\*\*\**P* < 0.001, n = 4). Error bars represent s.e.m. F. Hematoxylin and eosin (H&E) staining of testes in adult control and *Srsf10*^cKO^ mice. Scale bar, 50 μm. G. H&E staining of the cauda epididymis in adult control and *Srsf10*^cKO^ mice. Scale bar, 50 μm. H. Co-staining for MVH and PLZF (undifferentiated spermatogonia marker) in adult testes in control and *Srsf10*^cKO^ mice. DNA was stained with Hoechst 33342. Scale bar, 50 μm. I. Statistics of PLZF-positive cells per tubule of adult testes in control and *Srsf10*^cKO^ mice. At least 200 tubules were counted from at least three different mice. \*\*\**P* < 0.001. Error bars represent s.e.m.

The adult *Srsf10*^cKO^ males looked grossly normal. Normal copulatory plugs could be observed when adult *Srsf10*^cKO^ males were mated with wild-type females, but no pups were obtained (Table 1), indicating that *Srsf10*^cKO^ males were infertile. Compared to control, the testes of adult *Srsf10*^cKO^ males were much smaller (Figure 1C). The weight of testes and ratio of testes to body weight in *Srsf10*^cKO^ males were significantly reduced compared to the control (Figure 1D and 1E). We then analyzed the histology of adult *Srsf10*^cKO^ and control testes using hematoxylin and eosin (H&E) staining. While all populations of spermatogonia, spermatocytes and spermatids were observed in the control seminiferous tubules, fewer germ cells were observed and no spermatocytes and spermatids could be observed in the center of seminiferous tubules from adult *Srsf10*^cKO^ testes (Figure 1F). Moreover, no mature spermatozoa could be found in the cauda epididymis of *Srsf10*^cKO^ mice (Figure 1G). To further identity of the remaining germ cells in *Srsf10*^cKO^ testes, we co-stained these germ cells with MVH and PLZF (an undifferentiated spermatogonia-specific marker)^31^. Immunofluorescence results showed that only sporadic MVH^+^PLZF^+^ cells can be detected around the basement of seminiferous tubules from adult *Srsf10*^cKO^ testes (Figure 1H). Moreover, the number of PLZF^+^ cells was significantly reduced in adult *Srsf10*^cKO^ testes compared to the control (Figure 1I), suggesting that only a few undifferentiated spermatogonia were left in the adult *Srsf10*^cKO^ testes. Similarly, testes from one-month-old *Srsf10*^cKO^ were also much smaller and germ cells were absent (Figure 1- figure supplement 1). Taken together, these data demonstrate that *Srsf10* is essential for male fertility and spermatogenesis.

**Table 1:** The fertility of *Srsf10*^cKO^ males.

| Genotype |  | No. of male mice | No. of plugged female mice | No. of litters | No. of pups per litter |
| --- | --- | --- | --- | --- | --- |
| Male | Female |  |  |  |  |
| Control | Wt | 8 | 17 | 14 | 13.43±3.78 |
| *Srsf10*<sup>ckO</sup> | Wt | 8 | 16 | 0 | 0 |

### *Srsf10* depletion leads to failure of meiosis initiation

As no spermatocytes were observed in the seminiferous tubules of adult and one-month-old *Srsf10*^cKO^ testes, we speculated that the meiosis process failed in *Srsf10*^cKO^ mice. To figure out which stage in spermatogenesis was affected after *Srsf10* deletion, we carefully analyzed the sections from P8, P10, P12, and P15 testes using H&E staining^32^. At P8, Type A spermatogonia (A), intermediate spermatogonia (In) and type B spermatogonia (B) could be observed in both control and *Srsf10*^cKO^ testes. Afterward, some germ cells entered into meiosis and developed into leptotene spermatocytes in P10 control testes, and abundant zygotene and pachytene spermatocytes could be observed in P12 and P15 control testes, respectively (Figure 2A). However, rarely differentiated spermatocytes were detected in the seminiferous tubules of P10, P12 and P15 *Srsf10*^cKO^ testes. Instead, some apoptotic cells were observed in the seminiferous tubules of P10 and P12 *Srsf10*^cKO^ testes (Figure 2A), suggesting that meiosis initiation might be impaired in *srsf1*0 deleted mice.

**Fig. 2.**
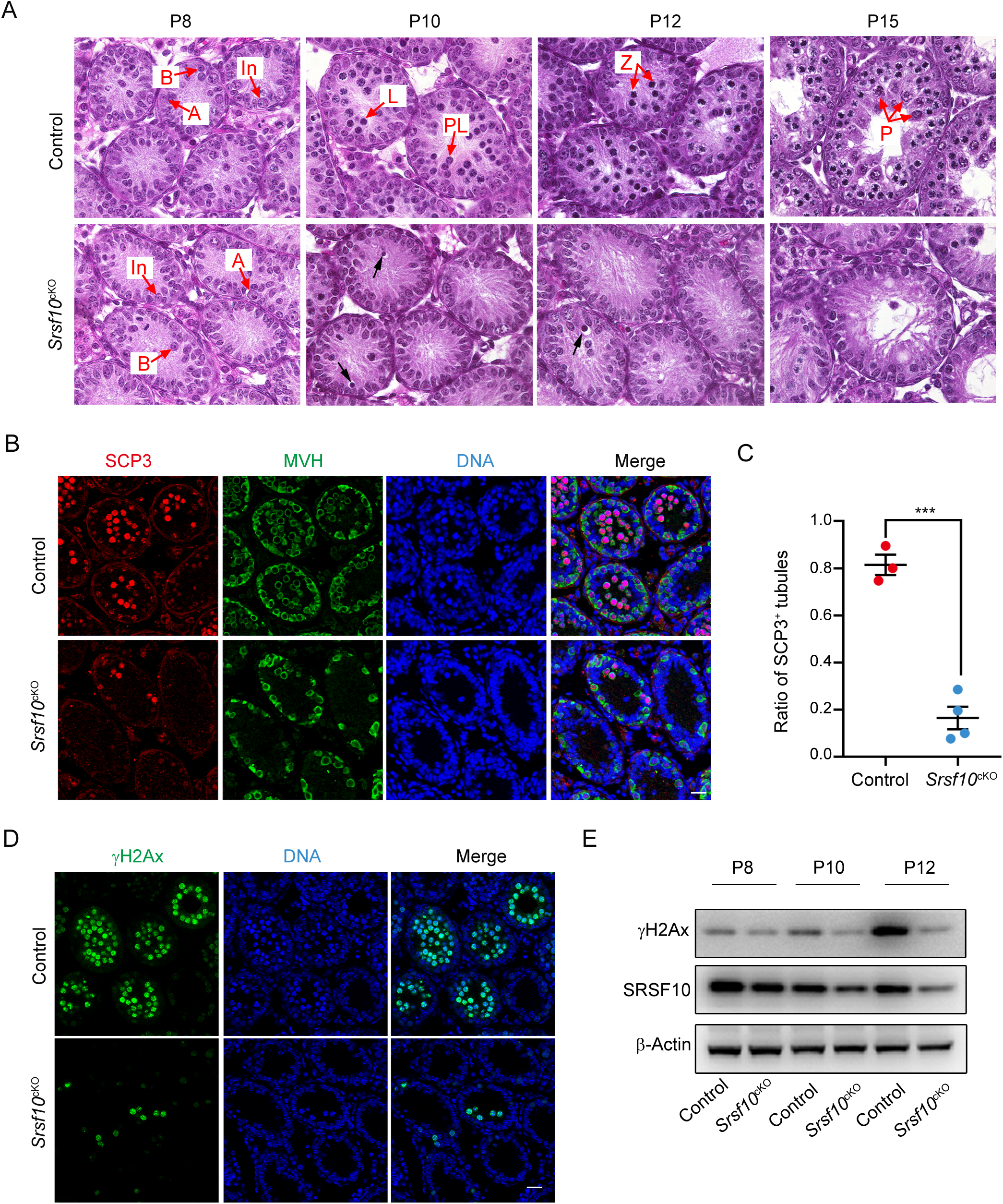
*Srsf10* deficient germ cells fail to initiate meiosis. A. H&E staining of control and *Srsf10*^cKO^ testes at P8, P10, P12 and P15. Spermatogenic cells were shown in cross-sections of seminiferous tubules from control and *Srsf10*^cKO^ testes. Scale bar, 10 μm. Red arrows indicate the representative stages of the spermatocytes. A, type A spermatogonia; In, intermediate spermatogonia; B, type B spermatogonia; L, leptotene spermatocytes; Z, zygotene spermatocytes; P, pachytene spermatocytes. Black arrows indicate apoptotic cells. Scale bar, 10 μm. B. Immunofluorescence co-staining for SCP3 and MVH in control and *Srsf10*^cKO^ testes at P12. Scale bar, 20 µm. C. Statistics of the ratio of SCP3-positive tubules in control and *Srsf10*^cKO^ testes at P12. At least 500 tubules were counted from at least three different mice. \*\*\**P* < 0.001. Error bars represent s.e.m. D. Immunofluorescence staining for γH2AX in control and *Srsf10*^cKO^ testes at P12. Scale bar, 20 µm. E. Western blot analyses of γH2AX in control and *Srsf10*^cKO^ testes at P8, P10 and P12. β-actin was used as the loading control.

To further confirm the above results, we co-stained MVH and SCP3, which was a component of the synaptonemal complex and the marker of meiosis prophase I at P12. Spermatocytes in the vast majority of seminiferous tubules had entered into meiosis prophase I as expected, and abundant MVH^+^SCP3^+^ germ cells could be detected in the center of control seminiferous tubules. However, only very few seminiferous tubules contained a tiny minority (about 3 to 5) of MVH^+^SCP3^+^ germ cells (Figure 2B). Statistically, the ratio of SCP3^+^ seminiferous tubules in *Srsf10*^cKO^ testes was significantly lower than the control at P12 (0.16 ± 0.048 versus 0.81 ± 0.043, mean ± SEM) (Figure 2C). Next, we detected the expression of the marker of homologous recombination γH2AX, which can be another indicator of meiosis in P12 testes. Immunofluorescence results showed that γH2AX^+^ cells were nearly undetectable in the seminiferous tubules of *Srsf10*^cKO^ testes (Figure 2D). Western blot showed that γH2AX was markedly elevated in control testes from P10 to P12, consistent with the entering into meiotic prophase I of many spermatocytes around this time. However, the expression of γH2AX changed little in *Srsf10*^cKO^ testes from P8 to P12 (Figure 2E), indicating failure of meiosis entrance. These results indicate that *Srsf10* deletion causes a severe defect in meiosis initiation during spermatogenesis.

### *Srsf10* depletion impairs the expansion and differentiation of the progenitor spermatogonia population

Before meiosis, SSCs proliferate to self-renew or generate differentiation-committed progenitors. The progenitor spermatogonia respond to the RA signal and divide into KIT^+^ differentiating spermatogonia (Figure 3A). Then type B differentiating spermatogonia further proliferate and differentiate into meiotic spermatocytes. Therefore, the initiation of meiosis requires several rounds of mitotic divisions for full expansion and efficient differentiation of spermatogonia^4, 33, 34^. Interestingly, we found that the expression of MVH and PLZF was significantly reduced in *Srsf10*^cKO^ testes from P8 to P12 (Figure 3B). Co-staining of MVH and PLZF showed that the number of PLZF^+^ undifferentiated spermatogonia was apparently reduced in *Srsf10*^cKO^ testes at P12 (Figure 3-figure supplement 1, A and B). These results indicate that the loss of *Srsf10* affects the normal development of spermatogonia, which may further lead to the failure of meiosis initiation.

**Fig. 3.**
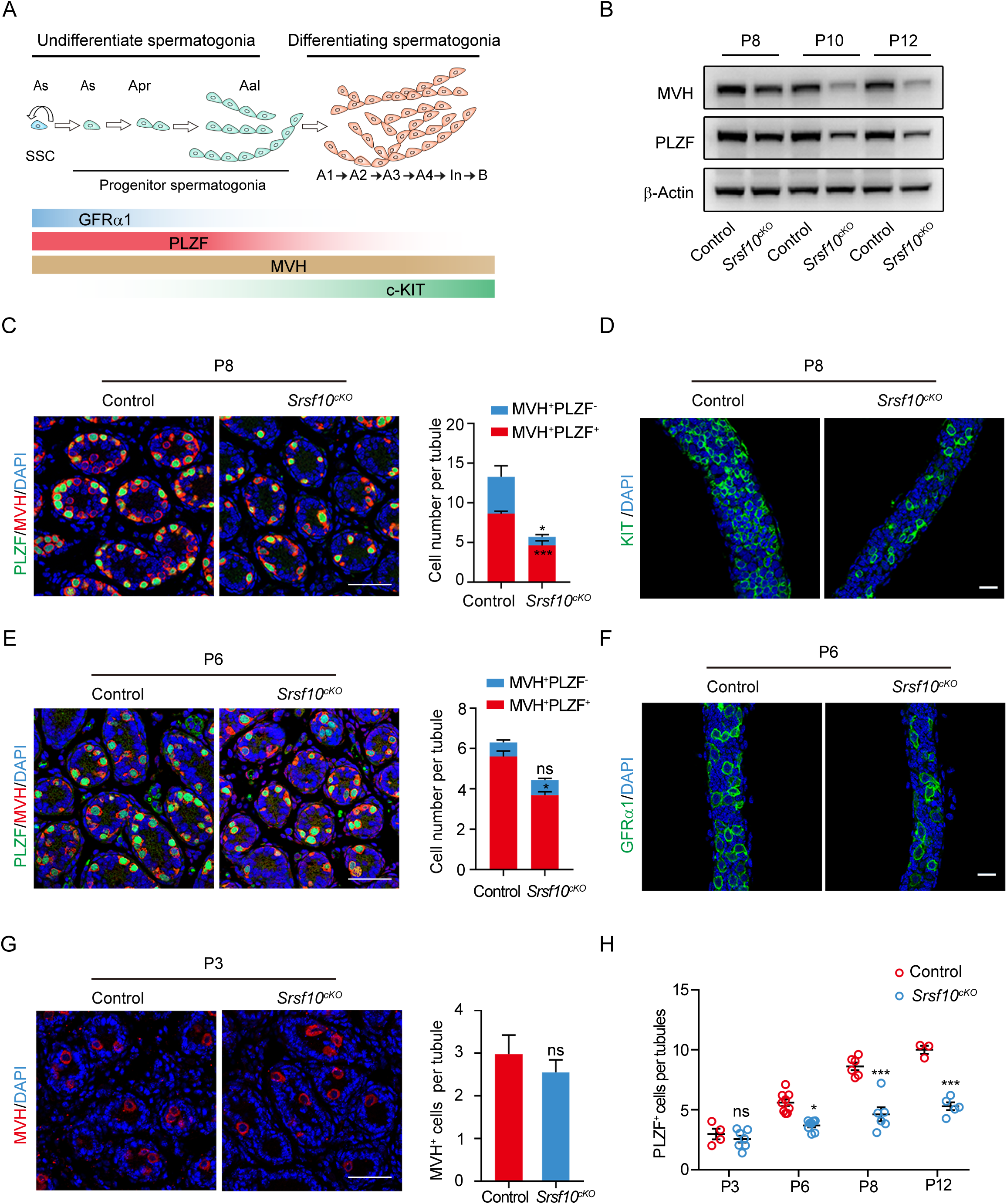
*Srsf10* depletion impairs the expansion and differentiation of progenitor spermatogonia. A. Schematic diagram showing the progression of mitosis phase of spermatogonia from spermatogonial stem cells (SSCs) to differentiating spermatogonia. The expression of representative markers at each stage is also shown. B. Western blot analyses of PLZF and MVH in control and *Srsf10*^cKO^ testes at P8, P10 and P12. β-actin was used as the loading control. C. Left, immunofluorescence co-staining for PLZF and MVH in control and *Srsf10*^cKO^ testes at P8. Scale bar, 50 µm. Right, quantification of MVH-positive and PLZF-positive cells (MVH^+^PLZF^+^) per tubule or MVH-positive but PLZF-negative (MVH^+^PLZF^+^) cells per tubule in control and *Srsf10*^cKO^ testes at P8. At least 1000 tubules were counted from 6 different mice. \*\*\**P* < 0.001. \**P* <0.05. Error bars represent s.e.m. D. Whole-mount staining for KIT in control and *Srsf10*^cKO^ testes at P8. Scale bar, 20 µm. E. Left, immunofluorescence co-staining for PLZF and MVH in control and *Srsf10*^cKO^ testes at P6. Scale bar, 50 µm. Right, quantification of MVH^+^PLZF^-^ cells or MVH^+^PLZF^+^ cells per tubule in control and *Srsf10*^cKO^ testes at P6. At least 1000 tubules were counted from at least 3 different mice. \**P* < 0.05. ns represents no significance. Error bars represent s.e.m. F. Whole-mount staining for GFRα1 in control and *Srsf10*^cKO^ testes at P6. Scale bar, 20 µm. G. Left, immunofluorescence staining for MVH in control and *Srsf10*^cKO^ testes at P3. Scale bar, 50 µm. Right, quantification of MVH-positive cells per tubule in control and *Srsf10*^cKO^ testes at P3. At least 1000 tubules were counted from at least 4 mice. ns represents no significance. Error bars represent s.e.m. H. Quantification of PLZF-positive cells per tubule in control and control and *Srsf10*^cKO^ testes at P3, P6, P8 and P12. P3 are counted with MVH-positive cells. \**P* < 0.05. \*\*\**P* < 0.001. ns represents no significance. Error bars represent s.e.m.

Considering the loss of PLZF^+^ cells in P12 *Srsf10*^cKO^ testes, we speculated that the proliferation and/or differentiation of spermatogonia may be disrupted when *Srsf10* is depleted. Therefore, we analyzed the distribution and the number of spermatogonia in both control and *Srsf10*^cKO^ testes at P8 and P6, when only spermatogonia and Sertoli cells can be observed in the seminiferous tubules. At P8, co-staining for MVH and PLZF showed that numerous differentiating spermatogonia (MVH^+^PLZF^-^) could be detected at the basement of seminiferous tubules in control. However, MVH and PLZF were nearly co-localized in *Srsf10*^cKO^ testes, indicative of few differentiating spermatogonia (MVH^+^PLZF^-^) (Figure 3C). Statistical analysis showed that differentiating spermatogonia (MVH^+^PLZF^-^) were nearly absent and the number of undifferentiated spermatogonia (MVH^+^PLZF^+^) was significantly reduced in *Srsf10*^cKO^ testes compared to the control (Figure 3C). We further performed whole-mount staining of the testes for the differentiating spermatogonia marker KIT.

The result showed that the KIT^+^ cells were obviously reduced in *Srsf10*^cKO^ testes compared to the control at P8 (Figure 3D), suggesting that deletion of *Srsf10* leads to inefficient differentiation of the progenitor spermatogonia. At P6, co-staining for MVH and PLZF showed that most of the germ cells were undifferentiated spermatogonia (MVH^+^PLZF^+^) in both control and *Srsf10*^cKO^ testes (Figure 3E). However, the number of the undifferentiated spermatogonia (MVH^+^PLZF^+^) was also significantly reduced in *Srsf10*^cKO^ testes compared to the control (Figure 3E). PLZF is a broader marker of all undifferentiated spermatogonia including As, Apr and Aal spermatogonia (Figure 3A). Thus, we performed whole-mount staining for GFRα1, which is specifically expressed in As and Apr spermatogonia^35, 36^, and showed that the GFRα1^+^ cells were also apparently reduced in *Srsf10*^cKO^ testes compared to the control at P6 (Figure 3F). Altogether, these results suggest that fewer undifferentiated spermatogonia (SSCs and progenitor cells) are produced in the absence of *Srsf10* as early as P6.

During male germ cell development, the pre-spermatogonia undergo a period of G1/G0 arrest from around E15.5 to P3, when these germ cells migrate from the center of seminiferous tubules to the periphery, and then restore mitosis proliferation, transforming into type A spermatogonia and some get stem cell properties around P3 to P6^37, 38^. Because *Vasa-Cre* recombinase is expressed as early as E15.5, we asked whether depletion of *Srsf10* would affect the development of pre-spermatogonia at P3. There was no difference in germ cell numbers between control and *Srsf10*^cKO^ testes (Figure 3G), suggesting that *Srsf10* is dispensable for the development of G1/G0 arrested pre-spermatogonia. Interestingly, from P3 to P12, the number of undifferentiated spermatogonia was gradually increased in both control and *Srsf10*^cKO^ testes. However, the kinetics in *Srsf10*^cKO^ testes was much slower, leading to a comparable number of PLZF^+^ cells per tubule in P12 *Srsf10*^cKO^ to that in P6 control testes (Figure 3H). Western blot confirmed that the expression of PLZF in P3 testes was lower and no significant difference was observed between control and *Srsf10*^cKO^ testes. With the resumption of mitosis, the expression of PLZF was apparently increased from P3 to P8 in control testes, but was only slightly increased in *Srsf10*^cKO^ testes (Figure 3-figure supplement 1C). It should be noted that we cannot exclude the possibility that the SSCs are also affected. However, the slower increase of PLZF^+^ cells suggests that *Srsf10* may be essential for the expansion and differentiation of the progenitor spermatogonia population.

### *Srsf10* depletion disturbs the expression of genes involved in progenitor spermatogonia

To systematically investigate the molecular effects of *Srsf10* loss in germ cells, we compared the transcriptomes of control and *Srsf10^cKO^* testes at P3, P6 and P8. RNA-seq analyses identified only 10 upregulated and 9 downregulated genes in *Srsf10^cKO^* testes compared with the control at P3 (FPKM >= 5, fold change >= 2, *P* < 0.01) (Figure 4A), suggesting that the transcriptome is largely normal at P3 in the *Srsf10* depletion testes. This is consistent with the immunostaining result showing that the number of germ cells is comparable between control and *Srsf10^cKO^* testes at P3. However, 139 genes were differentially expressed in the *Srsf10^cKO^* testes at P6 and nearly all genes (135/139) were down-regulated in *Srsf10^cKO^* testes (Figure 4A). 396 genes were differentially expressed in *Srsf10^cKO^* testes at P8, including 243 upregulated and 153 downregulated genes (Figure 4A). Surprisingly, only 57 genes were commonly downregulated in the *Srsf10^cKO^* testes at both P6 and P8 (Figure 4B). Gene ontology (GO) analysis of the 57 genes showed that these genes are involved in the meiotic cell cycle, spermatogenesis, cell differentiation and germ cell development (Figure 4B). We then systematically analyzed the expression pattern of all downregulated genes in control and *Srsf10^cKO^* testes from P3 to P8. The vast majority of down-regulated genes were gradually increased from P3 to P8 (Figure 4- figure supplement 1A). Interestingly, P6 specific down-regulated genes (e.g., *Pou5f1*, *GFRα1*, *Sohlh1*, *Egr4*, *Nanos3*, *Foxc2*, and *Sox3*) are mainly involved in the expansion and early differentiation of progenitor spermatogonia (Figure 4B). In P6 *Srsf10^cKO^* testes, the expression level of these genes was very low, although they were up-regulated afterward and reached a level comparable to the control P6 testes in P8 *Srsf10^cKO^* testes (Figure 4-figure supplement 1A). These results suggest that the expansion and early differentiation of progenitor spermatogonia are presumably impaired as early as P6, consistent with the observation that the number of PLZF^+^ and GFRα1^+^ undifferentiated spermatogonia was significantly reduced at P6 in the absence of *Srsf10*. Many P8-specific downregulated genes (e.g., *Kit*, *Sycp3*, *Sycp2*, *Atm* and *Rec8*) were involved in spermatogonia differentiation and meiosis and were barely up-regulated in *Srsf10^cKO^* testes from P6 to P8 (Figure 4B and Figure 4- figure supplement 1A), echoing nearly no differentiating spermatogonia and meiotic spermatocytes in P8 *Srsf10^cKO^* testes. Unsupervised cluster analysis of spermatogonia-specific genes (SPGs)^12^ showed that the gene expression pattern of *Srsf10^cKO^* testes at P8 was much more similar to that of control testes at P6 (Figure 4C), consistent with the compromised and slower expansion and differentiation of progenitor spermatogonia in the *Srsf10^cKO^* testes than control.

**Fig. 4.**
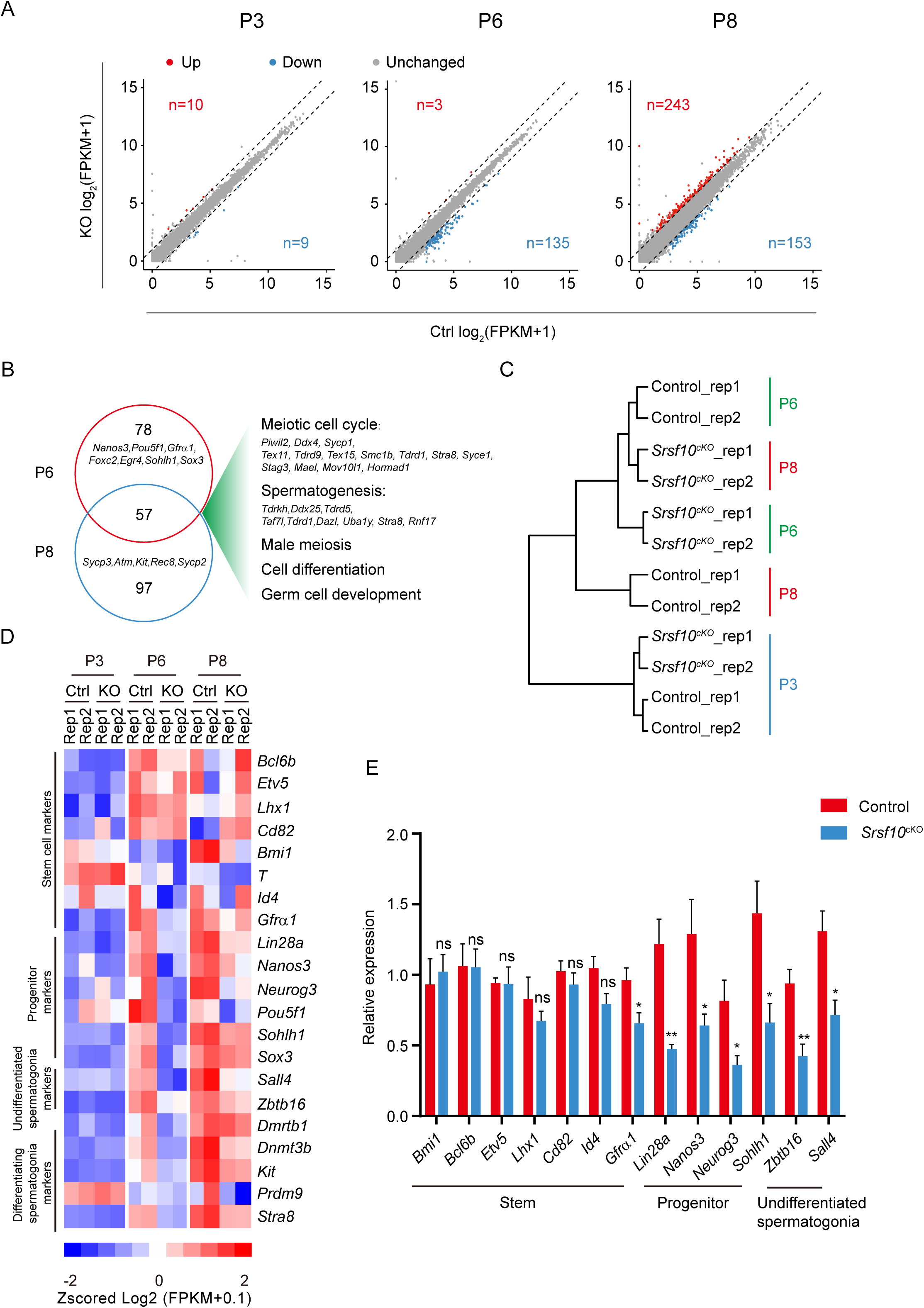
*Srsf10* deficiency alters expression patterns of genes involved in progenitor spermatogonia. A. Scatter plots showing the expression of genes in control and *Srsf10*^cKO^ testes at P3, P6 and P8. Blue dots represent significantly down-regulated genes, while red dots show significantly up-regulated genes (FPKM >= 5, fold change >= 2, *P* < 0.01). Grey dots represent unchanged genes. B. Venn diagram depicting the overlap of downregulated genes between P6 and P8. Gene ontology (GO) terms of the 57 shared downregulated genes in P6 and P8 *Srsf10*^cKO^ testes and genes involved in specific GO terms were shown on the right. C. Hierarchical clustering of two replicates of control and *Srsf10*^cKO^ testes at P3, P6 and P8 based on the expression of spermatogonia specific genes^12^. Note the closer relationship between P8 *Srsf10*^cKO^ and P6 control testes. D. Heatmap showing the mRNA abundance of genes functioning in SSCs (*Bcl6b*, *Etv5*, *Lhx1*, *Cd82*, *Bmi1*, *T*, *Id4* and *Gfra1*), progenitors (*Lin28a*, *Nanos3*, *Neurog3*, *Pou5f1*, *Sohlh1* and *Sox3*), undifferentiated spermatogonia (*Sall4* and *Zbtb16*) and differentiating spermatogonia (*Dmrtb1*, *Dnmt3b*, *Kit*, *Stra8* and *Prdm9*). E. Quantitative RT-PCR validation of the expression of genes involved in SSC, progenitors, undifferentiated spermatogonia and differentiating spermatogonia in control and *Srsf10*^cKO^ testes at P6. β-actin was used as the internal control. \**P* < 0.05, \*\**P* < 0.01, \*\*\**P* < 0.001; ns, no significance. Error bars represent s.e.m.

We then systematically analyzed several marker genes for stem cell maintenance, expansion and differentiation of the spermatogonia population. Genes involved in SSC maintenance (*Etv5*, *Bmi1*, *Id4*, *Lhx1*, *Cd82* and *T*) were hardly affected in the *Srsf10^cKO^* testes from P3 to P8. However, genes associated with progenitor spermatogonia (*Lin28a*, *Nanos3*, *Sox3*, *Neurog3*, *Pou5f1* and *Sohlh1*), undifferentiated spermatogonia (*Plzf* and *Sall4*) and differentiating spermatogonia (*Dmrtb1*, *Dnmt3b*, *Stra8*, *Kit* and *Prdm9*) were down-regulated in *Srsf10^cKO^* testes at P6 and P8 compared to control (Figure 4D and Figure 4- figure supplement 1B). The qRT-PCR results confirmed that *Srsf10* deficiency did not affect the expression of genes involved in SSCs, but globally reduced the expression of genes that control expansion and differentiation of progenitor cells at P6 (Figure 4E). These results are consistent with previous staining results that the number of stem/progenitor cells was significantly decreased at P6. Taken together, these data suggest that *Srsf10* deficiency may not affect the gene expression and formation of SSCs, but disturbs genes involved in progenitor spermatogonia and thus their expansion and differentiation.

### Single-cell RNA-seq reveals that the progenitor and differentiating spermatogonia are largely lost in *Srsf10*^cKO^ testes

To address the heterogeneity of spermatogonia after *Srsf10* depletion, we sought to enrich the undifferentiated and differentiating spermatogonia using their cell surface markers, THY1 (undifferentiated) and KIT (differentiating). We enriched THY1^+^ KIT^-^, THY1^-^KIT^+^ and THY1^+^ KIT^+^ spermatogonia from control and *Srsf10^cKO^* testes using magnetic-activated cell sorting (MACS) with anti-THY1 and anti-KIT antibodies at P8^39^. The isolated single cells were subjected to single-cell RNA-seq analysis using the 10X Genomics platform (Figure 5A). After filtering out low-quality cells and somatic cells, 1157 control and 766 *Srsf10^cKO^* (from two replicates) spermatogonial cells were used for further analysis. Uniform manifold approximation and projection (UMAP) and cell type-specific marker gene analyses were then performed for cell-type identification^40^. UMAP analysis identified five distinct spermatogonial subtypes/states that we named USSC1, USSC2, DSSC1, DSSC2 and DSSC3 (Figure 5B), mainly comprising undifferentiated spermatogonia and differentiating spermatogonia. Analysis using marker genes indicates that USSC1 cells correspond to SSCs, as they highly express SSC marker genes, including *Eomes, Id4, Lhx1, Etv5, Gfra1* and *Smoc2* (Figure 5C and Figure 5- figure supplement 1A). USSC2 cells are highly enriched for *Nanos3, Pou5f1* and *Plzf* (*Zbtb16*), with increased levels of *Sox3* and *Sohlh1*, but lower levels of SSC-associated markers and differentiation markers (Figure 5C and Figure 5- figure supplement 1A), corresponding to early progenitor spermatogonia. By contrast, DSSC1 cells express high levels of early differentiation marker genes such as *Stra8, Sox3, Sohlh1, Sohlh2* and *Dmrt1* and low levels of *Pou5f1, Nanos3* and *Plzf* (Figure 5C and Figure 5- figure supplement 1A), indicating that this cell population is likely late progenitor/early differentiating spermatogonia. DSSC2 cells display lower *Stra8* expression but mainly express differentiation marker genes including *Kit, Dmrtb1, Cited1* and *Dmrt1* (Figure 5C and Figure 5- figure supplement 1A), implicating them as differentiated spermatogonia. The DSSC3 cluster was only observed from the *Srsf10^cKO^* group. These cells express some differentiating markers such as *Stra8*, *Dmrt1*, Sohlh2 and *Kit*, but the expression level of *Stra8* and *Kit* was lower than DSSC1 and DSSC2, respectively (Figure 5C), suggesting that DSSC3 cells might have differentiated but with abnormal development. The pseudo-time analysis provided a trajectory indicating the development of germline cells from USSC1 to DSSC2 (Figure 5D).

**Fig. 5.**
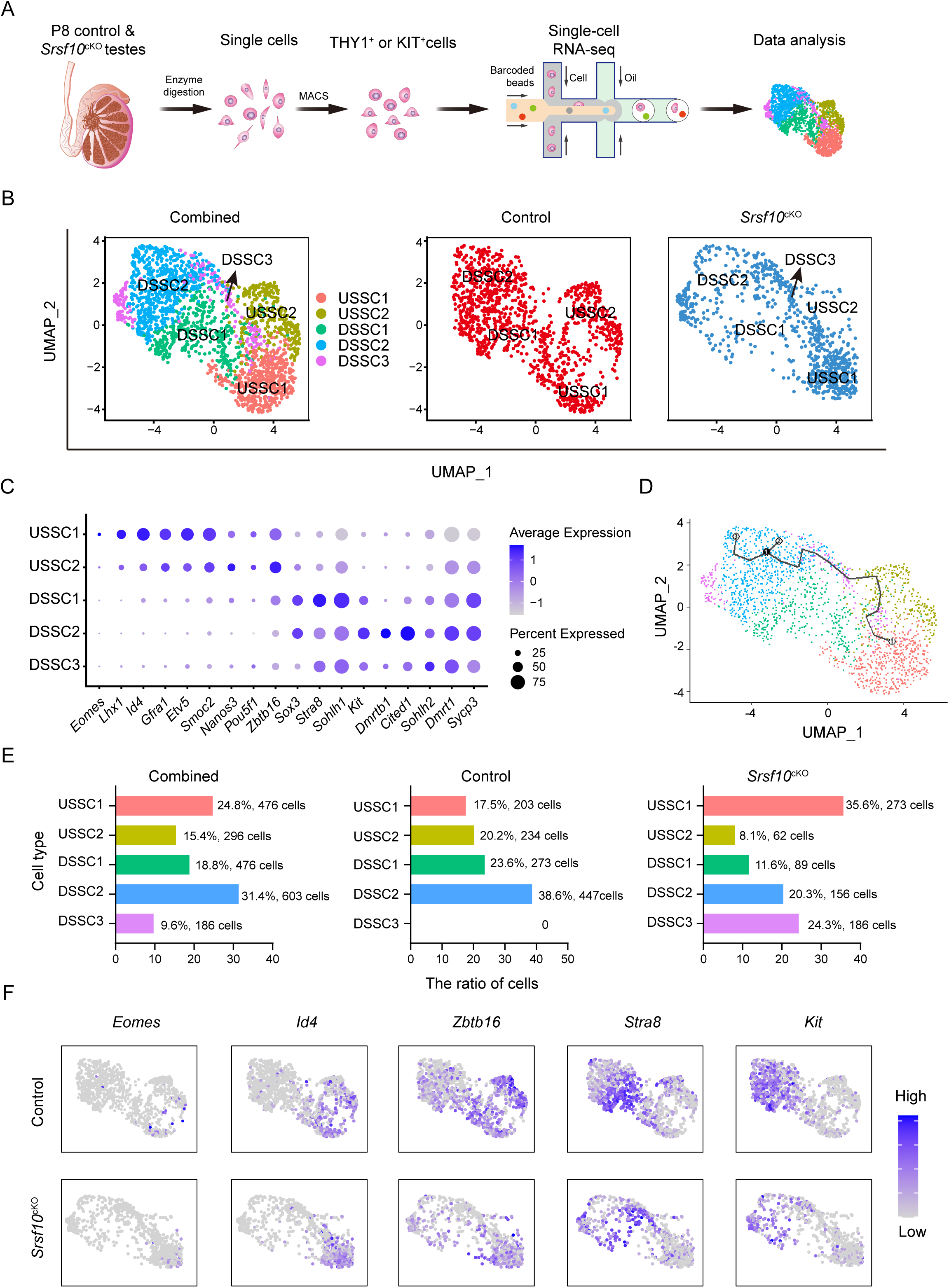
scRNA-seq defines the transcriptome-wide signatures of spermatogonia development in *Srsf10*^cKO^ testes at P8. A. Schematic illustration of the workflow for scRNA-seq analysis. B. UMAP clustering analysis of combined (left), control (middle), and *Srsf10*^cKO^ (right) spermatogonia. Five subtypes, including USSC1, USSC2, DSSC1, DSSC2 and DSSC3 are identified and color-coded in the left panel. C. Dot plot showing the expression of selected marker genes across the five spermatogonia subtypes. D. Pseudotime trajectory analysis of the indicated cell clusters. E. Quantification of the number and percentage of spermatogonia in each subtype in combined, control and *Srsf10*^cKO^ groups. F. The expression of selected marker genes in each subtype in control and *Srsf10*^cKO^ samples.

We then asked whether and how the subtypes of spermatogonia changed after *Srsf10* depletion. We found that USSC1, USSC2, DSSC1 and DSSC2 cells were largely evenly distributed in the control group, with 17.5% (203 cells), 20.2% (234 cells), 23.6% (273 cells) and 38.6% (447 cells) cells sorted into USSC1, USSC2, DSSC1 and DSSC2, respectively (Figure 5E). By contrast, the proportion in the *Srsf10^cKO^* group was severely slanted, which was resulted from the dramatic decrease of USSC2 (8.1%, 62 cells), DSSC1 (11.6%, 89 cells) and DSSC2 (20.3%, 156 cells) subtypes, concomitant with the dominance of the USSC1 subtype (35.6%, 273 cells). On the other hand, 24.3% (186 cells) of the cells were sorted into the DSSC3 subtype, which scattered in the other four spermatogonial subtypes (Figure 5E). This suggests that *Srsf10* deficiency severely impairs the development of spermatogonia. Analysis using SPG subtypes marker genes revealed that *Eomes^+^*, *Id4^+^* and *Gfra1^+^* cells (SSCs) were comparable in control and *Srsf10^cKO^* spermatogonia (Figure 5F and Figure 5- figure supplement 1B). *Zbtb16*^+^ and *Sall4*^+^ cells (undifferentiated spermatogonia), and *Nanos3*^+^ and *Egr4*^+^ cells (progenitor spermatogonia) were significantly reduced in the USSC2 subtype in the *Srsf10^cKO^* testes (Figure 5- figure supplement 1B). Early differentiating spermatogonia (*Stra8*^+^ and *Sohlh1*^+^ cells) and late differentiating spermatogonia (*Kit*^+^ and *Dmrtb1*^+^ cells) were also significantly reduced in the DSSC1 and DSSC2, respectively, in the *Srsf10^cKO^* testes (Figure 5F and Figure 5- figure supplement 1B). Taken together, single-cell transcriptional analysis of spermatogonia shows that the progenitor and differentiating spermatogonia are largely lost in *Srsf10^cKO^* testes at P8, further confirming the requirement of *Srsf10* for the expansion of progenitor spermatogonia

### The cell cycle and proliferation of spermatogonia are impaired in *Srsf10*^cKO^ testes

We then analyzed the differentially expressed genes in the aforementioned four subtypes of cells between the control and *Srsf10^cKO^* samples. In USSC1, 108 and 98 genes were down-regulated and up-regulated, respectively (|log2FC| > 0.25, adjusted *P*-value < 0.01) (Figure 6A). The down-regulated genes were mainly enriched for ‘Cell cycle’, ‘Cell division’, ‘Chromosome segregation’, ‘Spermatogenesis’ and ‘Regulation of G2/M transition of mitotic cell cycle’. The up-regulated genes were enriched for ‘G1/S transition of mitotic cell cycle’, ‘Regulation of apoptotic process’, ‘Regulation of cell proliferation’ and ‘Cellular response to growth factor stimulus’ (Figure 6B). Consistently, *Top2a*, *Mki67*, *Cdc20*, *Ccnb1*, *Ccna2*, *Cenpe*, *Rad50, Kif11, Nusap1* and *Prc1*, which are important for cell cycle and cell division, were significantly reduced in the USSC1 cells in the *Srsf10*^cKO^ sample (Figure 6B, 6C and Figure 6- figure supplement 1). *Dnaja1*, *Aspm*, *Ccnb1*, *Btg1*, *Racgap1*, *Sycp1*, *Sycp3* and *Mns1,* which are required in spermatogenesis, were also significantly reduced in the USSC1 cells in the absence of *Srsf10* (Figure 6B, 6C and Figure 6- figure supplement 1). In the USSC2, DSSC1 and DSSC2, the down-regulated genes were also enriched for ‘cell cycle’, ‘cell division’ and ‘cell proliferation’ (Figure 6- figure supplement 2). These data indicate that depletion of *Srsf10* in germ cells likely disrupts the normal cell cycle and proliferation of spermatogonia.

**Fig. 6.**
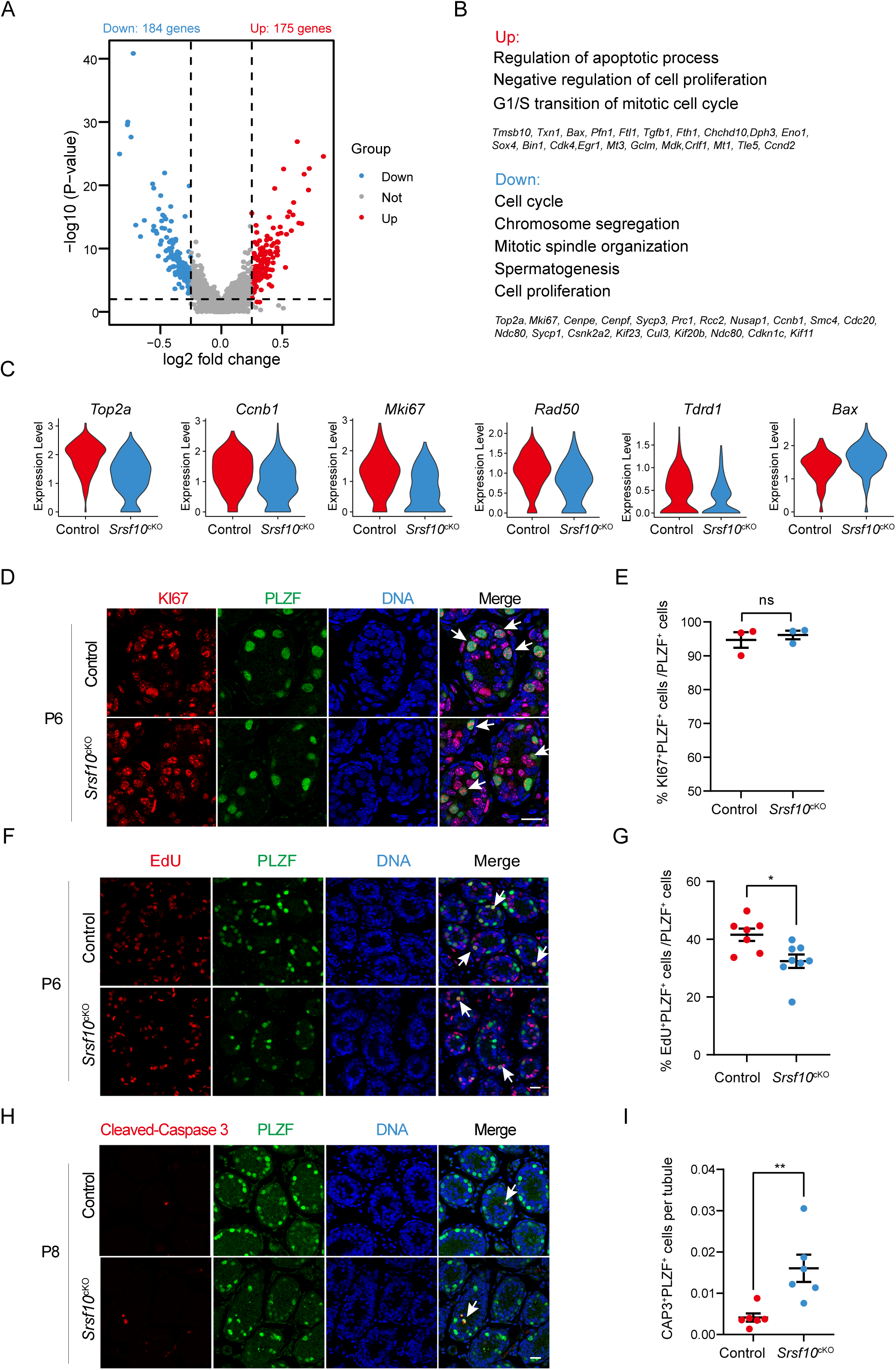
*Srsf10* depletion impairs the cell cycle and proliferation of undifferentiated spermatogonia. A. Volcano plot showing the significantly differentially expressed transcripts in the USSC1 subtype in *Srsf10*^cKO^ compared with the control samples. Blue dots represent significantly down-regulated transcripts, while red dots show significantly up-regulated transcripts (Log2 fold change >= 0.25, *P* < 0.01). Grey dots represent unchanged transcripts. B. Gene ontology of up-regulated and down-regulated genes in *Srsf10*^cKO^ USSC1 subtype and representative genes in up- and down-regulated groups are shown. C. Violin plots showing the expression of functional genes involved in cell cycle and spermatogenesis in the USSC1 subtype in control and *Srsf10*^cKO^ groups. D. Immunofluorescence co-staining for the mitosis marker KI67 and PLZF in control and *Srsf10*^cKO^ testes at P6. The DNA was stained with Hoechst 33342. Double-positive cells (KI67^+^PLZF^+^) are indicated by the white arrowhead. Scale bar, 50 μm. E. Quantification of the ratio of KI67^+^PLZF^+^ cells in PLZF^+^ cells in control and *Srsf10*^cKO^ testes at P6. ns, no significance. Error bars represent s.e.m. F. Immunofluorescence co-staining for the EdU and PLZF in control and *Srsf10*^cKO^ testes at P6. Control and *Srsf10*^cKO^ mice were treated with EdU for 4 hours. The DNA was stained with Hoechst 33342. White arrowheads indicate the representative EdU^+^PLZF^+^ cells. Scale bar, 20 μm. G. Quantification of the ratio of EdU^+^PLZF^+^ cells in PLZF^+^ cells of control and *Srsf10*^cKO^ testes at P6. At least 500 tubules were counted from at least 7 different mice. \**P* < 0.05, Error bars represent s.e.m. H. Immunofluorescence co-staining for the apoptosis marker cleaved caspase 3 (CAP3) and PLZF in control and *Srsf10*^cKO^ testes at P8. The DNA was stained with Hoechst 33342. Scale bar, 50 μm. I. Quantification of the number of CAP3^+^PLZF^+^ cells per tubule in control and *Srsf10*^cKO^ testes at P8. At least 300 tubules were counted from at least 5 different mice. \*\**P* < 0.01, Error bars represent s.e.m.

To test this idea, we first investigated the proliferation of spermatogonia in *Srsf10^cKO^* testes by co-staining for PLZF and KI67 (labeling mitotic cells that are not in the G0 phase) to analyze the mitotic status of spermatogonia at P6. The immunofluorescence result showed nearly all PLZF^+^ cells were also positive for KI67 in both control and *Srsf10^cKO^* testes (Figure 6D). The number of KI67^+^PLZF^+^ cells was not significantly different between control and *Srsf10^cKO^* testes (Figure 6E), suggesting that all the PLZF^+^ cells were in the mitotic status. We further tested the EdU incorporation of spermatogonia (PLZF^+^ cells) in the control and *Srsf10^cKO^* testes at P6. EdU is a thymidine analog and can incorporate into the DNA molecule during S-phase. Control and *Srsf10^cKO^* mice were intraperitoneally injected with EdU (5 mg/kg), and testes were collected 4 hours later for fixation and paraffin section. Co-staining for EdU and PLZF showed that the EdU incorporation was significantly reduced in *Srsf10^cKO^* PLZF^+^ spermatogonia at P6 (Figure 6F and 6G). These results suggest that the proliferation of spermatogonia in *Srsf10^cKO^* mice was significantly impaired. As apoptosis-related genes are up-regulated in the USSC1 and USSC2 cells, we also analyzed the apoptosis of spermatogonia. Double staining for cleaved caspase 3 (CAP3) and PLZF showed that the number of CAP3^+^PLZF^+^ spermatogonia was significantly higher in *Srsf10^cKO^* testes compared to the control (Figure 6H and 6I), implying that the survival of spermatogonia was also impaired when *Srsf10* is depleted. In conclusion, *Srsf10* depletion might lead to abnormal cell cycle and mitotic division, and further affect the proliferation and survival of spermatogonia.

### *Srsf10* regulates alternative splicing of functional genes in spermatogonia

We then sought to probe into how *Srsf10* depletion affects spermatogenesis at the molecular level. To this end, we collected the THY1^+^ spermatogonia using MACS from control and *Srsf10^cKO^* testes at P6 to minimize the potential side effects^39^ (Figure 7A). Flow cytometry and immunostaining analyses showed that the proportion of undifferentiated spermatogonia (THY1^+^ cells or PLZF^+^ cells) was significantly increased in the sorted group than that in the unsorted group (Figure 7- figure supplement 1A and 1B). We confirmed the dramatic decrease of *Srsf10* in the enriched spermatogonia of *Srsf10^cKO^* testes at both RNA and protein levels (Figure 7- figure supplement 1C and 1D). Then, we performed next-generation sequencing of the enriched spermatogonia of control and *Srsf10^cKO^* and compared the transcriptome between them. A total of 507 genes were differentially expressed in P6 *Srsf10^cKO^* spermatogonia (*P* < 0.01, fold change >= 2 and FPKM >= 2), including 200 down-regulated genes that were significantly enriched for genes involved in spermatogenesis, meiotic cell cycle, cell differentiation, male germline stem cell asymmetric division and germ cell development (Figure 7- figure supplement 2). Among the 200 down-regulated genes were the markers of progenitor spermatogonia, such as *Nanos3, Sox3, Lin28a* and *Neurog3* (Figure 7- figure supplement 2), presumably due to the absence of the progenitor spermatogonia, as observed in the scRNA-seq data.

**Fig. 7.**
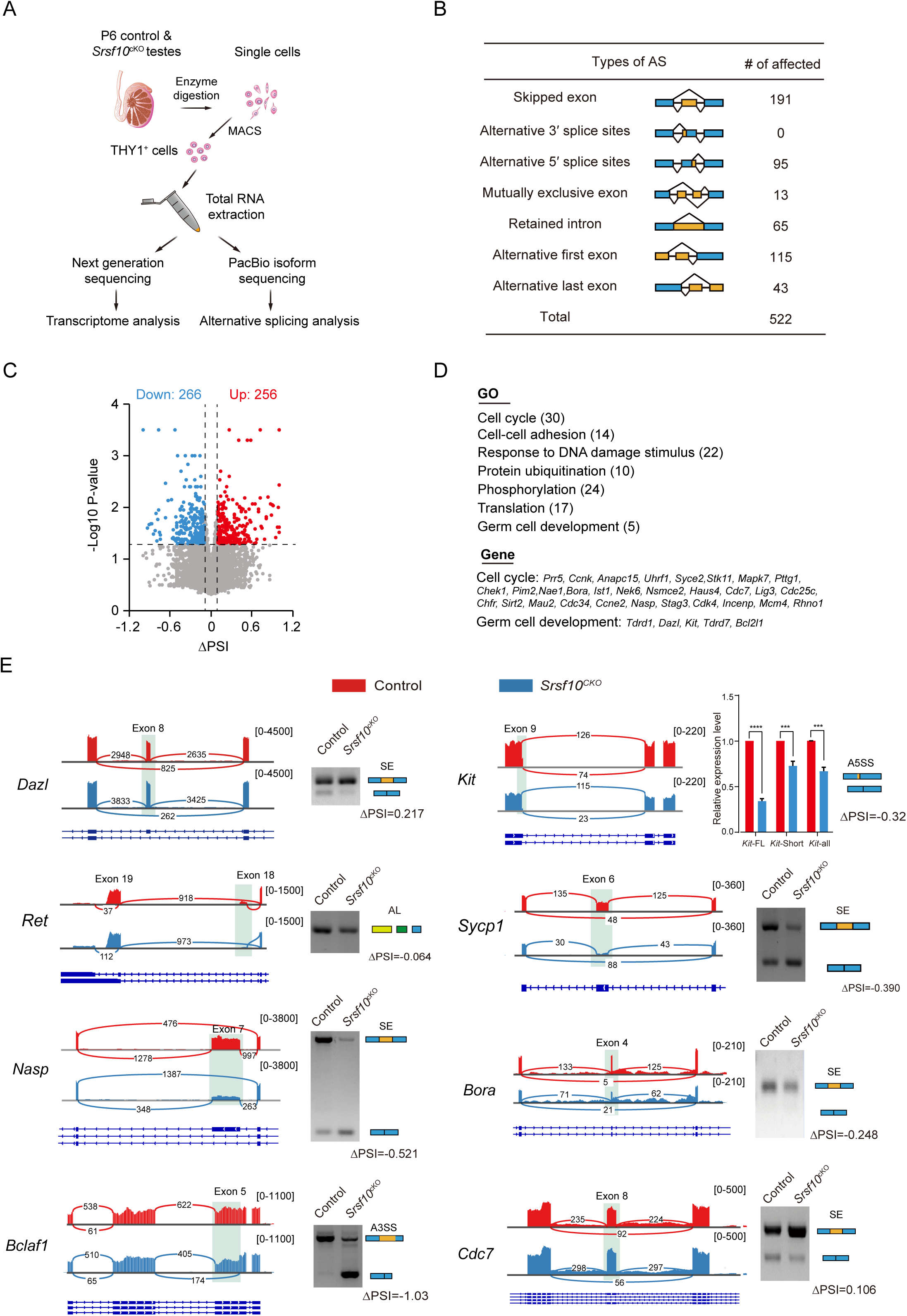
*Srsf10* is required for normal splicing of key genes involved in the cell cycle and germ cell development. A. Schematic illustration of the workflow for the enrichment of THY1^+^ spermatogonia for transcriptome analysis and alternative splicing analysis using Iso-seq. B. Seven AS events were significantly affected by depletion of *Srsf10* in the spermatogonia at P6. Splicing events affected by depletion of *Srsf10* were analyzed using SUPPA2 software (*P* <0.05 and |ΔPSI| >= 0.1). C. Scatter plot showing the significantly affected AS events (color-coded) in the absence of *Srsf10* at P6. Blue dots represent significantly down-regulated AS events (ΔPSI <= -0.1), while red dots show significantly up-regulated AS events (ΔPSI >= 0.1). D. Gene ontology terms for genes with significantly affected AS events and representative genes involved in cell cycle and spermatogenesis are shown. E. Visualization and validation of the differentially spliced genes in control and *Srsf10* depleted spermatogonia. Tracks from IGV are shown for selected candidate genes (left). Differentially spliced exons are shaded. Schematics of alternative splicing events are shown (blue and yellow rectangles) (right). Changes in “percent spliced in (PSI)” between control and *Srsf10*^cKO^ spermatogonia are shown below splicing schematics (ΔPSI). SE, Skipped exon, A5SS, Alternative 5’ splice sites, AL, Alternative last exon, A3SS, Alternative 3’ splice sites.

*Srsf10* is an SR protein and is involved in constitutive and alternative splicing. We asked whether depletion of *Srsf10* would affect the splicing and alternative splicing in spermatogonia. THY1^+^ spermatogonia at P6 were collected and subjected to PacBio Isoform Sequencing (Iso-seq) (Figure 7A), which can generate and capture the full-length cDNAs and present more accurate information about isoforms, alternatively spliced exons and fused transcripts. We then analyzed the Iso-seq data for identification of the differential alternative splicing events between control and *Srsf10^cKO^* spermatogonia using SUPPA2 software^41^. 522 differentially spliced events (DSEs) were identified (P <0.05 and |ΔPSI| > 0.1), the majority of which were exon skipping (191) (Figure 7B). Among the 522 DSE, 266 AS events had negative ΔPSI values and 256 had positive ΔPSI values, indicating that *Srsf10* has no bias in the regulation of AS (Figure 7C). The 522 DSEs involve 419 genes. Gene ontology (GO) analysis revealed that these genes participate in cell cycle (30 genes), cell-cell adhesion (14 genes), response to DNA damage stimulus (22 genes) and germ cell development (5 genes) (Figure 7D). We then asked what would be resulted from these DSEs. Taking exon skipping as an example, 132 of the 191 differentially spliced exons (excluded or included) happens in the gene coding regions (Figure 7- figure supplement 3A). While 59.8 % (n=79) of them produces another protein-coding isoform of potential regulatory functions (such as, *Dazl*, *Nasp* and *Cdc7*) (Figure 7- figure supplement 3B), about 40% of them generates frame-shifted transcripts (n=53) (Figure 7- figure supplement 3A). Notably, the latter can potentially give rise to the premature termination codon (PTC), such as in *Bora*, *Kat7* and *Clk1* (Figure 7- figure supplement 3C). The presence of PTCs probably further results in nonsense-mediated mRNA decay (NMD). Moreover, the next-generation sequencing (NGS) data were analyzed for alternative splicing using CASH software which is an alternative splicing event detecting method with a high rate of validation^42^. 332 differentially spliced events (DSEs) involved in 317 genes were identified (*P* <0.05) and the majority of which were exon skipping (229) (Figure 7-figure supplement 4A). 59 genes were commonly detected in the NGS data for CASH analysis and Iso-seq data for SUPPA2 analysis (Figure 7- figure supplement 4B). Gene ontology (GO) analysis of the 59 genes showed that these genes are involved in DNA replication (*Kat7*, *Mcm4*, *Nasp* and *Ssrp1*), cell cycle (*Bora*, *Cdc7*, *Stag3*, *Mcm4* and *Uhrf1*) and spermatid development (*Xlr5a*, *Xlr5b* and *Xlr5c*) (Figure 7- figure supplement 4B). In addition to these overlapped genes, we also focused on the functional splicing changes involved in spermatogenesis and cell cycle. for example, *Ret*, *Sycp1*, *Exo1*, *Zfp207*, *Ccna2* and *Ola1* in CASH analysis data and *Ccne2*, *Kit*, *Ist1*, *Mapk7*, *Pttg1* and *Clk1* in SUPPA2 analysis data (Figure 7- figure supplement 4B).

We first confirmed the aberrant splicing of *Bclaf1* (skipped exon), *Acly* (skipped exon) and *Zfp207* (skipped exon) (Figure 7E and Figure 7- figure supplement 4C), which are known targets of *Srsf10*^27, 43^. Importantly, the differential splicing of individual events in genes involved in important spermatogonia-related processes, including cell cycle and germ cell development were also successfully verified. For example, loss of *Srsf10* resulted in differential splicing of exon 8 in *Dazl* (increased exon inclusion), alternative last exon of *Ret*, alternative 5’ splice sites of exon 9 in *Kit* (absence of four amino acids), exon 6 in *Sycp1* (increased exon skipping), exon 8 in *Cdc7* (increased exon inclusion), exon 7 in *Nasp* (increased exon skipping), exon 4 in *Bora* (increased exon skipping), exon 14 in *Kat7* (increased exon skipping), exon 7 in *Ccna* (increased exon skipping), exon 9 in *Ist1* (increased exon skipping), alternative 5’ splice sites of exon 10 in *Exo1* and alternative first exon of *Ccne2* (Figure 7E and Figure 7- figure supplement 4C). These data suggest that SRSF10 is required for the correct splicing of functional genes, thus maintaining the homeostasis of alternative splicing during spermatogenesis.

## Discussion

Spermatogenesis is a very complex process in which germ cells undergo mitosis division of spermatogonia, genomic rearrangement in meiosis, and morphological changes in spermiogenesis to give rise to millions of mature spermatozoa per day^44^. The foundation of this process is spermatogonial stem cells (SSCs) which can self-renew to maintain the stem cell pool or undergo differentiation into spermatogonial progenitors that expand and further differentiate. Therefore, fully understanding the regulation of proliferation and differentiation of SSCs is of great importance. Recently, the importance of alternative splicing as a post-transcriptional regulatory mechanism involved in spermatogonia development is just beginning to be unraveled^13, 45^. However, the knowledge is still very limited regarding how the splicing machinery, including specific splicing factors that are involved in this process. By illustrating the function and mechanism of *Srsf10* in mouse spermatogenesis (Figure 8), our study highlights the complexity and requirements of alternative splicing regulation in spermatogonia expansion and meiosis initiation.

**Fig. 8.**
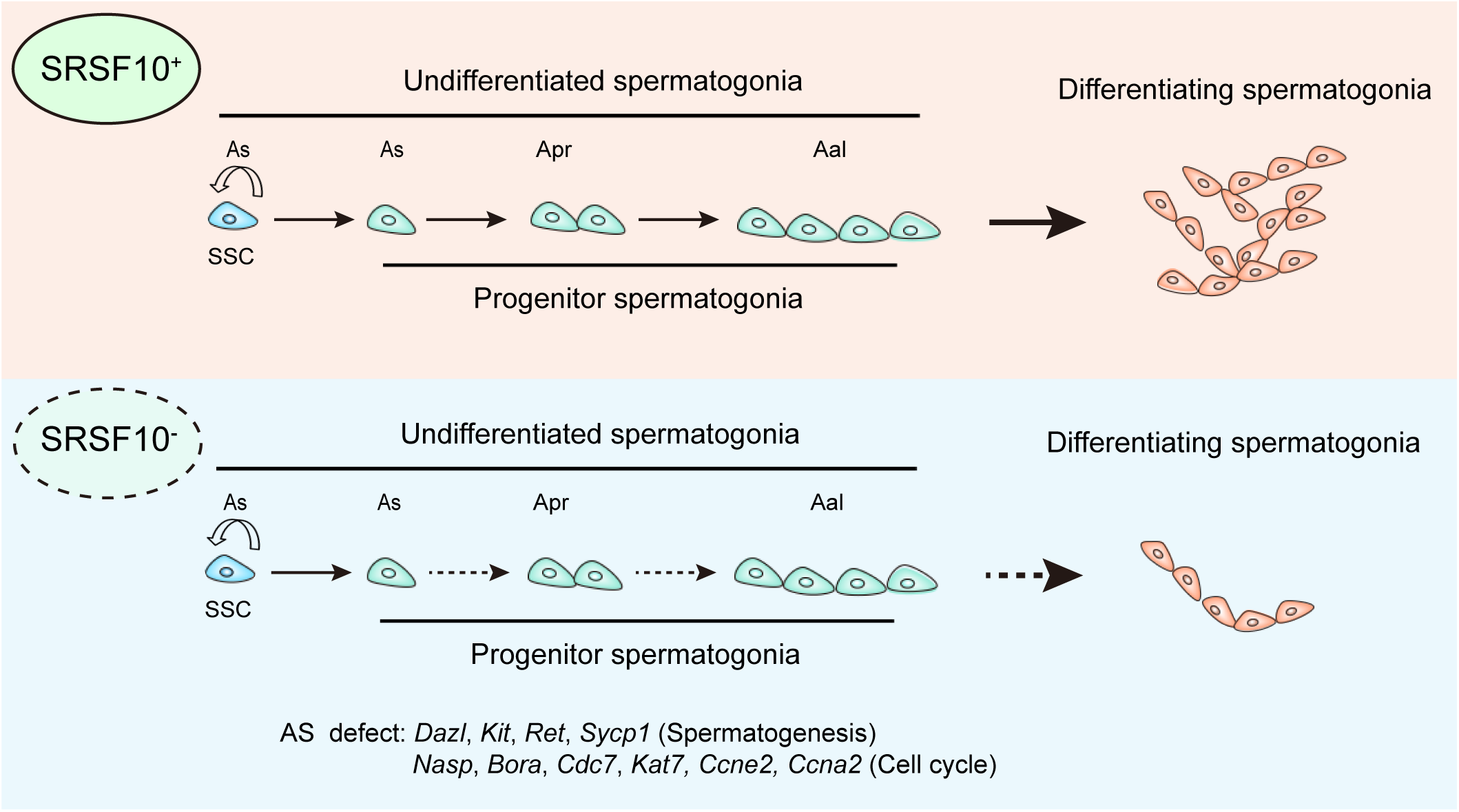
Model of SRSF10-mediated AS regulation in the development of undifferentiated spermatogonia. In the control testes, SRSF10 promotes the expansion of progenitor spermatogonia and subsequent differentiation of spermatogonia. In the absence of SRSF10, the alternative splicing (AS) defects of key genes involved in spermatogenesis and cell cycle impair the expansion of progenitor spermatogonia, leading to the decreased number of undifferentiated spermatogonia and nearly absence of differentiating spermatogonia.

*Srsf10* depletion mediated by Vasa-Cre leads to absence of differentiating spermatogonia, with existence of only a few undifferentiated spermatogonia in adult testes, and thus male infertility. In the first wave of spermatogenesis, the number of PLZF^+^ undifferentiated spermatogonia increases but with a much slower pace from P6 to P12, and differentiating spermatogonia and spermatocytes can barely be observed. This phenotype is very different from that of the depletion of genes that are important for the maintenance of SSCs (*Etv5*, *Bcl6b*, *Id4* and *Taf4b*)^39, 46, 47, 48^ or differentiation of spermatogonia (*Sohlh1* and *Sohlh2*)^49, 50^. The former does not lead to infertility until adulthood but exhibits an age-dependent progressive loss of germ cells, while the latter leads to loss of differentiating spermatogonia but with limited effect on undifferentiated spermatogonia. So, it is likely that the defects of *Srsf10^cKO^* testes might mainly be due to failed expansion of the progenitor spermatogonia population, and thus their efficient differentiation. This is further supported by the RNA-seq data of P3, P6 and P8 testes and scRNA-seq data of P8 spermatogonia, which show that SSCs can form but progenitor cells are reduced. So *Srsf10* is possibly critical for the formation and expansion of progenitor spermatogonia, but not the formation or maintenance of SSCs.

scRNA-seq data showed that the gene expression associated cell cycle of SSCs and progenitor cells was abnormal in *Srsf10^cKO^* testes. This is further supported by the reduced EdU incorporation in PLZF^+^ spermatogonia in the *Srsf10^cKO^* testes as early as P6. The PLZF^+^ spermatogonia include SSCs and progenitor cells and the proliferation of progenitor spermatogonia is reportedly at a higher rate than SSCs which are normally quiescent and exhibit a very slow cell cycle^51^. The scRNA-seq data show that the subtypes of SSCs can form and the number of SSCs is comparable to control. Thus, the phenotype of significantly decreased undifferentiated spermatogonia is most likely due to the abnormal cell cycle or proliferation of progenitor cells. Interestingly, alternative splicing shows that ‘cell cycle’ sits atop the *Srsf10* affected gene GO terms and contains the most genes (30, including *Prr5, Ccnk, Anapc15, Uhrf1, Syce2, Stk11, Mapk7, Pttg1, Check1, Pim2, Nae1, Bora, Ist1, Cdc7, Cdc25c, Ccne2, Nasp, Stag3, Cdk4* and *Mcm4,* etc), many of which have been successfully verified. Therefore, the negative effects on cell proliferation of spermatogonia of *Srsf10* depletion might be due to abnormal alternative splicing of abundant functional genes (Figure 8). However, how the aberrant splicing of these genes affects their functions and further leads to the phenotype needs to be further clarified.

Interestingly, besides the above factors that are involved in general cellular events, we found and verified the aberrant splicing of many important germ cell genes, including *Dazl*, *Kit*, *Ret* and *Sycp1* in *Srsf10* deficient spermatogonia (Fig. 8). *Dazl* encodes an RNA-binding protein essential for germ cell development and meiotic initiation^52^. DAZL has been proved to promote the expansion and differentiation of progenitor spermatogonia^53, 54^. In our previous study, we found that *Bcas2* depletion in spermatogonia leads to dramatic skipping of exon 8 (decreased level of *Dazl*-FL isoform and increased level of Dazl-Δ8 isoform), which might further lead to a decreased level of total DAZL at the protein level^18^. However, *Srsf10* loss in spermatogonia results in increased inclusion of exon 8 (decreased level of *Dazl*-Δ8 isoform and increased level of *Dazl*-FL isoform). The roles of *Dazl*-FL and *Dazl*-Δ8 isoforms in spermatogenesis are unclear yet, but the specific target mRNAs for both *Dazl* isoforms were identified in ESCs, implying the potential functional divergence of these two isoforms in spermatogenesis^55^. Although how the increased inclusion of exon 8 in *Dazl* affects its function and spermatogenesis remains to be further clarified, we speculate that the splicing of *Dazl* should be precisely regulated to keep the balance of its two isoforms and exert the normal functions. KIT is the marker of differentiating spermatogonia and is critical for the survival and proliferation of differentiating type A spermatogonia^56^. Several isoforms of *c-Kit* are produced by alternative mRNA splicing, of which two are the presence or absence of four amino acids (GNNK+ and GNNK-, respectively) in the extracellular domain^57^. In the *Srsf10^cKO^* testes, the GNNK+ isoform was more decreased than GNNK-. Thus, the ratio of GNNK-/GNNK+ displayed a significantly elevated in the *Srsf10^cKO^* testes. Previous studies showed that the two KIT isoforms share a similar affinity to ligand SCF, but the expression of GNNK-isoform was stronger transformation activity in NIH3T3 cells and more tumorigenic in nude mice^58, 59^. Thus, the abnormal ratio of *Kit* isoforms may play a role in the development of spermatogonia. Glial cell-line-derived neurotrophic factor (GDNF) is critical for SSCs self-renewal in mice by binding to the GFRA1/RET receptor^60^. GDNF-mediated RET signaling is required the fate of undifferentiated spermatogonia and *Ret* mutant testis shows severe SSC depletion by P7 during the first wave of spermatogenesis^61^. What’s more, the numbers of progenitor spermatogonia decrease more severely than the numbers of SSCs when GDNF/RET signaling is inhibited^62^. Thus, we suggest the AS change of *Ret* regulated by *Srsf10* may also play important role in the spermatogonia development.

While it is widely accepted that alternative splicing is an important post-transcriptional regulation, its function in the specific context and biological processes has just begun to be illustrated. Our results demonstrate that SRSF10 is essential for the expansion and differentiation of progenitor spermatogonia by regulating alternative splicing (Figure 8), providing new insights into the regulation of spermatogenesis regarding how alternative splicing is involved by specific splicing factor.

## Materials and Methods

### Mice

All mice were maintained under specific-pathogen-free (SPF) conditions and the illumination time, temperature and humidity were all in accord with the guidelines of the Animal Care and Use Committee of the Institute of Zoology at the Chinese Academy of Sciences (CAS). The *Srsf10*^Floxed/Floxed^ (*Srsf10*^F/F^) mice were purchased from Shanghai Model Organisms Center, Inc. The *Srsf10*^F/F^ male mice were mated with *Vasa-Cre* transgenic females to obtain the mouse model with *Srsf10* specific deletion in the germ cell lines. All the mice had C57BL/6J genomic background. Genotyping of *Srsf10* was performed by PCR of mice tail genomic DNA. Forward primer: 5-AACATTTAGCACATTTGAGGAT-3, and reverse primer: 5-AACAGCCATATTAACCCGTCTTG-3 were used to detect the wild-type allele (615 bp) and the floxed allele (467 bp). Forward primer: 5- AGCATGCCTATCTTGTGT-3, and reverse primer: 5-TAACCCGTCTTGTAGTAAATCT-3 were used to detect the mutant allele (300 bp). The *Vasa-Cre* was genotyped with forward primer (5- CACGTGCAGCCGTTTAAGCCGCGT-3) and reverse primer (5-TTCCCATTCTAAACAACACCCTGAA-3) with the product was 240 bp. The annealing temperature of all primers was 58 °C. Four genotypes in the progeny, including *Srsf10*^F/+^, *Srsf10*^F/-^, *Srsf10^F/+^;Vasa-Cre* and *Srsf10^F/-^;Vasa-Cre* were identified. The genotype of *Srsf10^F/-^;Vasa-Cre* was used as mutants and was referred to as *Srsf10*^cKO^. The genotypes of *Srsf10^F/+^;Vasa-Cre* were used as control.

### Histological analysis, immunostaining and imaging

For histological analysis, testes from control and *Srsf10*^cKO^ were isolated and fixed in Bouin’s solution (Saturated picric acid: 37% Formaldehyde: Glacial acetic acid= 15: 5: 1) overnight at room temperature. These testes were dehydration through a graded ethanol (30%, 50%, 70%, 80%, 90%, 100% and 100%) and embedded in paraffin. Then, paraffin-embedded samples were cut into sections of 5-μm thickness. After dewaxing and hydration, the sections were stained with hematoxylin and 1% eosin and imaged with a Nikon ECLIPSE Ti microscope.

For immunostaining, testes from control and *Srsf10*^cKO^ mice were isolated and fixed in 4% PFA overnight at 4 °C. Following dehydration, the testes were embedded in paraffin and cut into sections of 5 μm thickness. After dewaxing and hydration, the sections were boiled in citrate antigen retrieval solution (0.01 M citric acid/sodium citrate, pH 6.0) for 20 mins in the microwave oven. After free cooling, the sections were washed with PBS 3 times and blocked with 5% BSA in PBS for 1 h at room temperature. Then, the sections were incubated with primary antibody overnight at 4 °C. On the second day, we washed the sections with PBS 3 times and incubated them with secondary antibody for 1 h at room temperature.

After washing in PBS 3 times, the sections were incubated with 2 mg/ ml of Hoechst 33342 (Sigma, B2261) for 15 min at room temperature. Finally, the sections were washed with PBS 2 times and mounted with Fluoromount-G medium (Southern Biotech, 0100-01). The immunofluorescence staining was imaged with a laser scanning confocal microscope LSM880 (Carl Zeiss, Germany).

The primary antibodies used were listed as follows, rabbit anti-SRSF10 polyclonal antibody (ab254935, Abcam, 1:200); goat anti-PLZF antibody (AF2944, R&D, 1:200); goat anti-KIT antibody (AF1356, R&D, 1:200); goat anti-GFRα1 antibody (AF560, R&D , 1:200); rabbit anti-DDX4/MVH polyclonal antibody (ab13840, Abcam, 1:200); rabbit anti-phospho-Histone H2A.X (Ser139/Tyr142) Antibody (#5438, Cell Signaling Technology, 1:200); mouse anti-SCP3 antibody (ab97672, Abcam, 1:200); rabbit anti-Cleaved Caspase-3 (Asp175) antibody (#9661, Cell Signaling Technology, 1:200); and rabbit anti-Ki67 antibody (ab15580, Abcam, 1:200).

The secondary antibodies used were listed as follows: Alexa Fluor 488 donkey anti-rabbit (Jackson, 1:500); Alexa Fluor 549 donkey anti-rabbit (Jackson, 1:500); Alexa Fluor 488 donkey Anti-Mouse (Jackson, 1:500); Alexa Fluor 549 donkey anti-Mouse (Jackson, 1:500); and Alexa Fluor 488 donkey Anti-goat (Jackson, 1:500).

### RNA extraction and qRT-PCR

Total RNA was extracted from whole testes or enriched cells using RNAzol® RT (Molecular Research Center. Inc, RN 190) following the manufacturer’s instructions. After removing the residual genomic DNA with the DNase I Kit (Promega, M6101), 500 ng of total RNA was reverse-transcribed into cDNAs using the PrimeScript RT Reagent Kit (TaKaRa, RR037A) according to the manufacturer’s protocol. qRT-PCR was performed using an Eva Green 2X qPCR MasterMix-No Dye kit (Abm, MasterMix-S) on a LightCycler 480 instrument (Roche). Relative gene expression was analyzed based on the 2^-ΔΔCt^ method with *β-actin* as internal controls. At least three independent experiments were analyzed. All primers were listed in the Supplementary file 1.

### Western blot

The protein from testes or enriched cells were extracted using RIPA lysis buffer (50 mM Tris–HCl (pH 7.5), 150 mM NaCl, 1% Sodium deoxycholate, 1% Triton X-100, 0.1% SDS, 5 mM EDTA, 1 mM Na3VO4, 5–10 mM NaF) containing a protease inhibitor cocktail (Roche, 04693132001) and the protein lysis buffers were incubated on ice for 20 min. After the ultrasound, the protein lysis buffers were centrifuged at 4 °C, 12,000 rpm for 20 min and quantified using a BCA reagent kit (Beyotime, P0012-1). After being boiled at 95 °C for 6 min, equal amounts of total protein lysates were used for immunoblotting analysis. Different molecular weight proteins were separated in a 10% SDS–PAGE gel and transferred onto PVDF membranes. After blocking with 5% non-fat milk for 1 h at room temperature, the membranes were incubated with diluted primary antibodies at 4 °C overnight. After three washes with TBST, the membranes were incubated with secondary antibodies conjugated with horseradish peroxidase (1:3,000, Jackson ImmunoResearch) at room temperature for 1 h. The signals were developed with Pierce ECL Substrate (Thermo Fisher Scientific, #34080), detected with Bio-RAD ChemiDocTMXRs+ and analyzed with Quantity One software (Bio-Rad Laboratories). All primary antibodies were listed as follows, rabbit anti-SRSF10 polyclonal antibody (NB110-93598, Novus Biologicals, 1:1,000); goat anti-PLZF antibody (AF2944, R&D, 1:1,000); rabbit anti-DDX4/MVH polyclonal antibody (ab13840, Abcam, 1:1,000); rabbit anti-phospho-Histone H2A.X (Ser139/Tyr142) Antibody (#5438, Cell Signaling Technology, 1:1,000).

### Whole-mount immunostaining

The testes were collected and dissected to remove the tunica albuginea. Seminiferous tubules were dispersed with tweezers and fixed at 4% PFA overnight at 4 °C. The tubules were washed three times with PBST for 10 min each and permeated with 0.1% TritonX-100 for 4 h at room temperature. Then, the tubules were washed three times with PBST and blocked with 5% BSA in PBS for 2 h at room temperature. Primary antibodies were incubated overnight at 4 °C. After washing with PBST and the tubules were incubated with secondary antibodies for 4 h at room temperature. With the washing of PBST, the tubules were stained with Hoechst for 30min at room temperature. After simply washing, the tubules were mounted with Fluoromount-G medium (Southern Biotech, 0100-01). The imaging of whole-mount staining followed the protocol described above.

### RNA sequencing

Testes samples were collected from P3, P6 and P8 control and *Srf10*^cKO^ mice. THY1^+^ spermatogonia were isolated from P6 control and *Srf10*^cKO^ mice. Total RNA was isolated using the RNAzol® RT (Molecular Research Center. Inc, RN 190) according to the manufacturer’s protocol and treated with DNase I to remove residual genomic DNA. A total amount of 1 μg of RNA per sample was used to prepare cDNA libraries generated using the NEBNext Ultra RNA Library Prep Kit for Illumina (NEB) following the manufacturer’s recommendations. 6G base pairs (raw data) were generated by Illumina Novaseq 6000 for each cDNA library. The adaptor sequence and sequences with a high content of unknown bases or low-quality reads were removed to produce the clean reads used for bioinformatic analysis.

### RNA-seq bioinformatic analyses

After initial quality control, the clean reads were aligned to the mouse reference genome (mm 9) using Tophat v2.1.1 (TopHat: discovering splice junctions with RNA-Seq). Next, the gene expression was calculated by Cufflinks v2.2.1. And the normalization of the gene expression values was based on the fragments per kilobase of exon per million reads mapped (FPKM).

### EdU incorporation assay

EdU (RiboBio, C00053) dissolved in PBS was injected intraperitoneally at 5 mg/kg of body weight. Testes were collected from control and *Srsf10*^cKO^ mice at 4 h following EdU incorporation and fixed in 4% PFA overnight at 4 °C, and then embedded in paraffin. EdU and PLZF co-staining was performed according to the protocol of Cell-Light^TM^ Apollo567 Stain Kit (100T) (RiboBio, C10310-1).

### Testes digestion and generation of cell suspensions

Testes from P6 mice were used to generate single-cell suspensions following enzymatic digestion as described previously^2^. In brief, testes were collected from P6 or P8 control and *Srsf10*^cKO^ mice in Hanks’ balanced salt solution (HBSS) and the tunica albuginea was removed. Then, the testes were digested with 4.5 ml 0.25% trypsin/EDTA and 0.5 ml 7 mg/ml DNase I solution (Sigma, d5025) for 6 min at 37 °C. After gently pipetting, another 0.5 ml DNase I solution was added to the cell suspension to digest for another 2-3 min at 37 °C and followed by the addition of 1 ml 10% FBS. Single-cell suspensions were made by gently repeated pipetting and passed through a 40 μm pore size cell strainer. The cells are centrifugated at 600 g for 7 min at 4 °C and resuspended with 180 μl running buffer (Miltenyi Biotec, 130-091-221) for MACS selection.

### THY1^+^ spermatogonia isolation

We added 20 μl THY1 antibody-conjugated microbeads [anti-mouse CD90.2 (Thy1.2) MicroBeads; Miltenyi Biotech, 130-121-278] to the single-cell suspensions and mixed well, then incubated for 20 min at 4 °C. After incubation, 2 ml running buffer was added to the cells and centrifugated at 600 g for 7 min at 4 °C. The pellet was then resuspended in 500 μl of running buffer for magnetic cell separation. The LS columns (Miltenyi Biotec, 130-042-401) were placed in the magnetic field of the MACS Separator and prewashed with 0.5 ml of running buffer followed by the addition of the THY1-labeled cell suspension. Wash the LS columns 3 times with 3 ml running buffer and the unlabeled THY1^-^ cells were eluted. Then, the LS columns were removed from the separator and placed on new 15 ml centrifuge tubes. The THY1^+^ cells were eluted from the LS columns in a 5 ml running buffer and centrifuged at 600g for 7 min at 4 °C. After centrifugation, the supernatant is removed and the pellet of THY1^+^ cells was stored at -80 °C for subsequent sequencing analysis and validation.

### Droplet-based single-cell RNA-sequencing

THY1^+^ KIT^-^, KIT^+^ THY1^-^ and THY1^+^KIT^+^ spermatogonia collection: Testes collected from P8 control and *Srsf10*^cKO^ mice were digested and the single cell suspensions were generated following the protocol described above. We added 20 μl THY1 antibody-conjugated microbeads [anti-mouse CD90 (Thy1.2) MicroBeads; Miltenyi Biotech, 130-121-278] and 40 μl KIT antibody-conjugated microbeads [anti-mouse CD117 MicroBeads; Miltenyi Biotech, 130-091-224] to the single-cell suspensions for 20 min at 4 °C, and followed the protocol described above to get the THY1^+^ KIT^-^, KIT^+^ THY1^-^ and THY1^+^KIT^+^ spermatogonia.

Single cells were suspended in PBS containing 0.04% BSA and loaded onto the Chromium 3’ v3 platform (10x Genomics) to generate single cell libraries according to the manufacturer’s protocol. Briefly, single cells were partitioned into Gel Bead-In-EMulsions (GEMs) in the 10X Chromium Controller instrument followed by cell lysis and barcoded reverse transcription using a unique molecular identifier (UMI). The cDNA was generated and then amplified, and the libraries were finally sequenced using an Illumina Novaseq 6000 sequencer with a sequencing depth of at least 100,000 reads per cell with a pair-end 150 bp (PE150) reading strategy (performed by CapitalBio Technology, Beijing).

### Data processing of scRNA-seq

Raw data were processed using the Cell Ranger Software (version 4.0.0). Briefly, the raw files were converted to demultiplexed fastq files through the Cell Ranger mkfastq pipeline. The fastq files were then aligned to the mouse reference genome (mm10) using the STAR aligner. Next, the reads were further filtered for valid cell barcodes and UMIs to produce a count matrix. Finally, the count matrix was imported into the R package Seurat and quality control was performed to remove outlier cells and genes. Cells with 200–4000 detected genes were retained. Genes were retained in the data if they were expressed in ≥ 3 cells. After applying these quality control criteria, 1923 cells and 23310 genes remained for further analysis. Additional normalization was performed in Seurat on the filtered matrix to obtain the normalized count. Highly variable genes across single cells were identified and PCA was performed to reduce the dimensionality on the top 16 principal components. Then, cells were clustered and visualized in two dimensions using UMAP (Uniform Manifold Approximation and Projection).

### Pseudotime analysis of single-cell transcriptomes

Germ cell lineage trajectories were constructed according to the procedure recommended in the Monocle3 start, cluster 2 and 3 (USSC2 and DSSC1) as middle, and cluster 5 (DSSC3) as the end of pseudotime. Briefly, documentation (https://cole-trapnell-lab.github.io/monocle3/docs/starting/). Cluster 1 (USSC1) was used as the top differentially expressed genes were selected as “ordering genes” to recover lineage trajectories in Monocle3 using default parameters. After pseudotime time was determined, differentially expressed genes were clustered to verify the fidelity of lineage trajectories.

### PacBio Isoform Sequencing (Iso-seq)

THY1^+^ spermatogonia were isolated from P6 control and *Srf10*^cKO^ mice. Total RNA was isolated using the RNAzol® RT (Molecular Research Center. Inc, RN 190) according to the manufacturer’s protocol and treated with DNase I to remove residual genomic DNA. 5 μg of total RNA per sample was used to prepare the Iso-Seq library using the Clontech SMARTer cDNA synthesis kit and the BluePippin Size Selection System protocol as described by Pacific Biosciences (PN 100-092-800-03).

After running the Iso-Seq pipeline, sequence data were processed using the SMRTlink 5.0 software. A circular consensus sequence (CCS) was generated from subread BAM files. Additional nucleotide errors in consensus reads were corrected using the Illumina RNAseq data with the software LoRDEC. Then all the consensus reads were aligned to the mouse reference genome (mm 9) using Hisat2.

### Differential splicing analysis and validation

Differential splicing events of Iso-seq data were analyzed using SUPPA2 software with the standard protocol. The SUPPA2 software can identify the seven common modes of AS events and obtain accurate splicing change quantification between control and *Srsf10*^cKO^ samples. We used *P*-value < 0.05 and |ΔPSI | > 0.1 as the threshold to filter for significantly differential splicing events. Differential splicing events of NGS data were analyzed using CASH software with the standard protocol using *P*-value < 0.05 as the threshold for filtering. For validation, we first imported the differentially spliced sites of interesting functional genes analyzed by SUPPA2 and CASH into the integrative genomics viewer tool (IGV) to efficiently and flexibly visualize and explore spliced sites between control and *Srsf10*^cKO^ samples. The primers for differentially spliced exons were designed using Primer5. Primers were designed within constitutive exons flanking the differentially spliced exons. Standard PCR for analysis by gel electrophoresis was performed according to the manufacturers’ instructions and visualized by running on a 2% agarose gel. All primers were listed in the Supplementary file 2.

### Statistical analysis

All experiments were performed at least three independent times. At least three independent biological samples were collected for the quantitative experiments. Quantification of positively stained cells was counted from at least three independent fields of view. Paired two-tailed Student’s t-test was used for statistical analysis and data were presented as mean ± SEM. *P*-value < 0.05 was considered with a significant difference level. Equal variances were not formally tested. No statistical method was used to predetermine sample sizes.

## Data availability

All RNA-sequencing data have been deposited to GEO with the accession number GSE190646. The following secure token has been created to allow review of record GSE190646 while it remains in private status: gbuvgeiqxjclbgf. Our work did not generate any datasets or use any previously published datasets. Source data for Figures 1, 2, 3, 4, 6, 7 and figure supplement of Figures 1, 3, 7 have been provided.

## Author contributions

W.L. designed and performed the major experiments and wrote the manuscript. K.L. performed the data analysis of RNA-seq, AS analysis and the drawing of a pattern diagram. Z.Z. contributed to the data analysis of single-cell RNA sequencing. Q.L., Z.G., Y.X. S.S. and W.L. contributed to mouse maintenance and performed some experiments. L.L., A.G. and H.L. contributed to some statistics and analysis. Z.H., Y.H., and Y. O. contributed to technical assistance. J.L., Q.S. and Z.W. analyzed the data and revised the manuscript. All authors read and approved the final manuscript.

## Acknowledgments

We thank Shiwen Li and Xili Zhu for their technical assistance. Jian Chen and all members in Sun lab for their helpful advice. We thank Mingming Fan for her guidance of MACS of spermatogonia.

---

**Figure 1-figure supplement 1.**
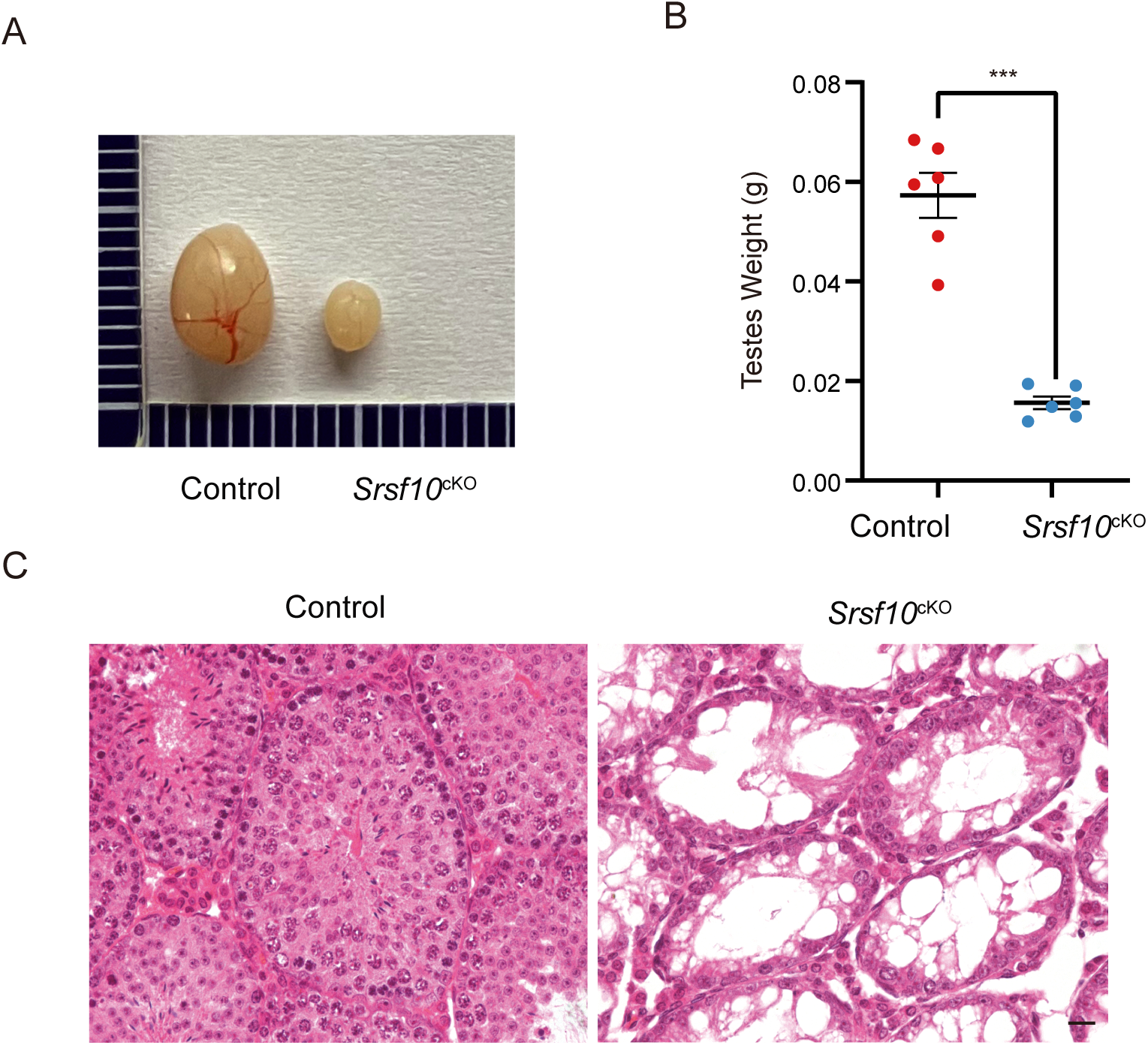
*Srsf10* is required for the first wave of spermatogenesis. A. Morphological analysis of the control and *Srsf10*^cKO^testes at P35. B. Testes weight of P35 control and *Srsf10*^cKO^ mice (\*\*\**P* < 0.001, n = 3). Error bars represent s.e.m. C. Hematoxylin and eosin (H&E) staining of P35 testes in control and *Srsf10*^cKO^ mice. Scale bar, 10 μm.

**Figure 1-source data 1** The testis weight and the number of PLZF^+^ cells of adult male mice.

**Figure 1- figure supplement 1-source data 1** The testis weight of one-month-old male mice.

**Figure 2-source data 1** The ratio of SCP3^+^ tubules in P12 testes.

**Figure 3-figure supplement 1.**
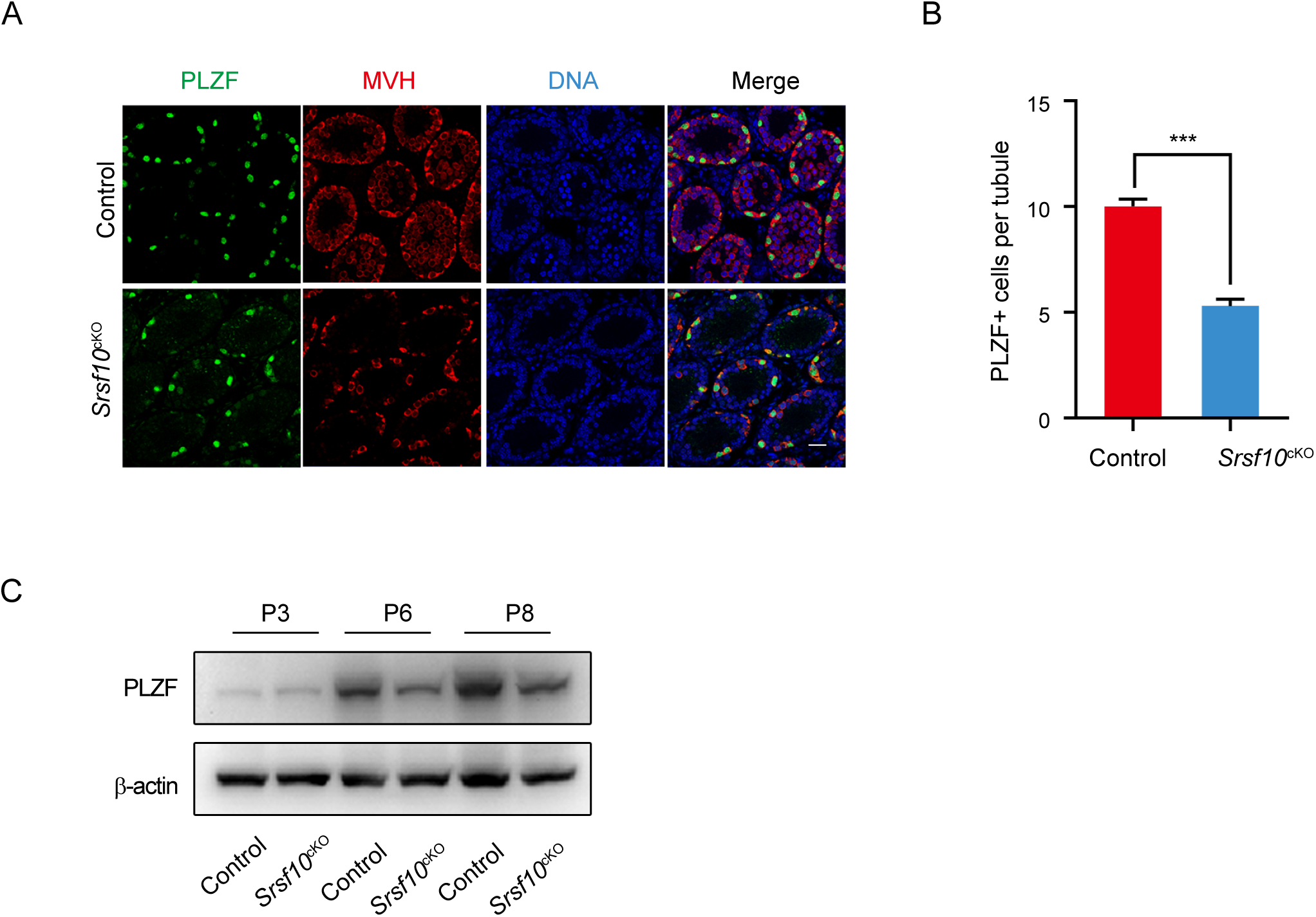
Spermatogonia development is impaired in *Srsf10*^cKO^ testes. A. Immunofluorescence co-staining for PLZF and MVH in control and *Srsf10*^cKO^ testes at P12. Scale bar, 20 µm. B. Quantification of PLZF-positive cells per tubule in control and *Srsf10*^cKO^ testes at P12. At least 300 tubules were counted from at least 3 different mice. \*\*\**P* < 0.001, Error bars represent s.e.m. C. Western blot analyses of MVH and PLZF in control and *Srsf10*^cKO^ testes at P3, P6 and P8. β-actin was used as the loading control.

**Figure 3-source data 1** Quantification of MVH/PLZF-positive cells per tubule in control and *Srsf10*^cKO^ testes at P3, P6, P8 and P12.

**Figure 3- figure supplement 1-source data 1** Quantification of PLZF-positive cells per tubule in P12 control and *Srsf10*^cKO^ testes.

**Figure 4-figure supplement 1.**
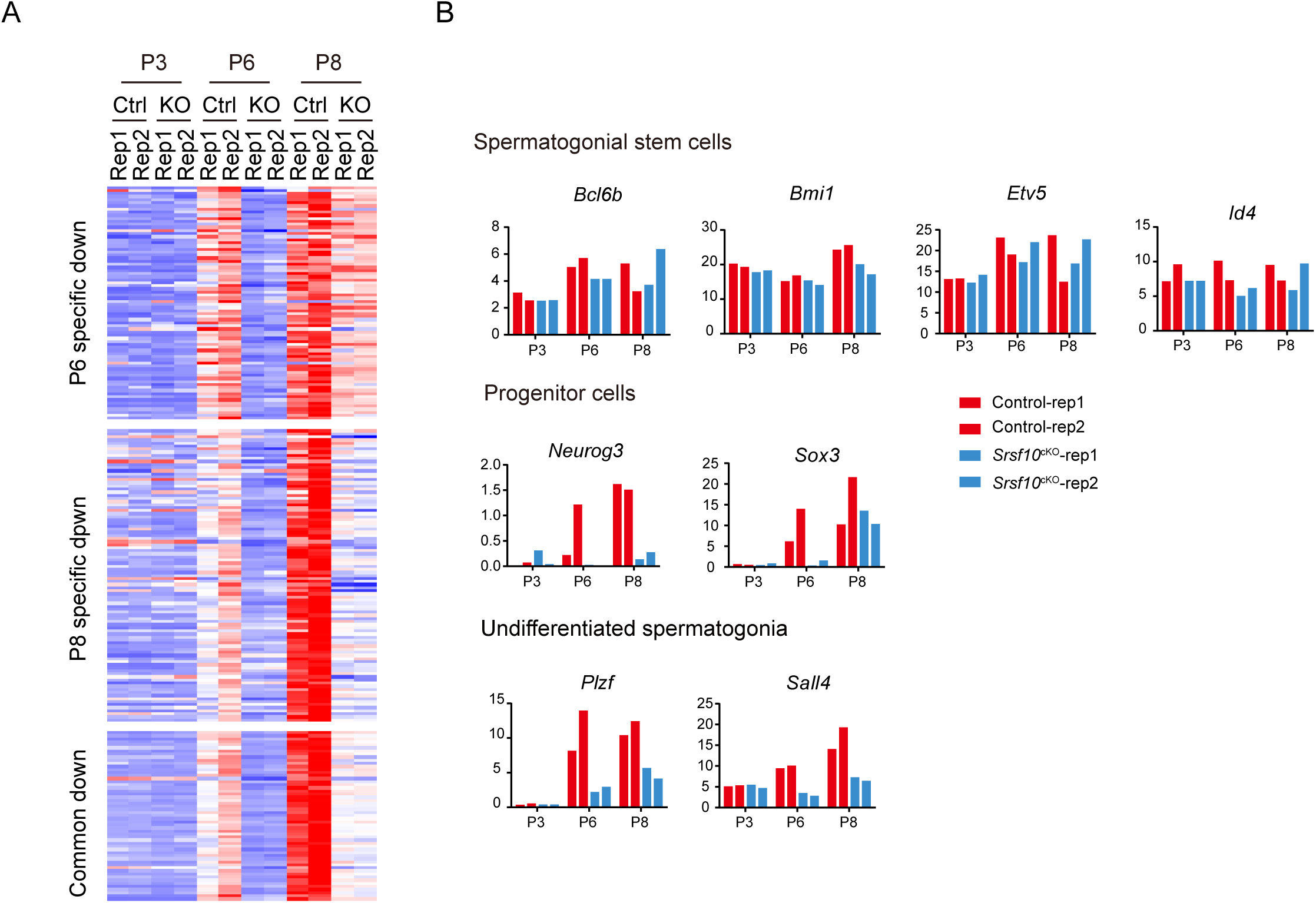
Systematical analysis of the down-regulated genes from P3 to P8. A. Heatmap shows the mRNA abundance of all the down-regulated genes from P3 to P8. B. RNA-seq results of the expression level of selected marker genes related to spermatogonial stem cells, progenitor cells and undifferentiated spermatogonia from P3 to P8.

**Figure 4-source data 1** The expression of marker genes in control and *Srsf10*^cKO^ testes at P6.

**Figure 5-figure supplement 1.**
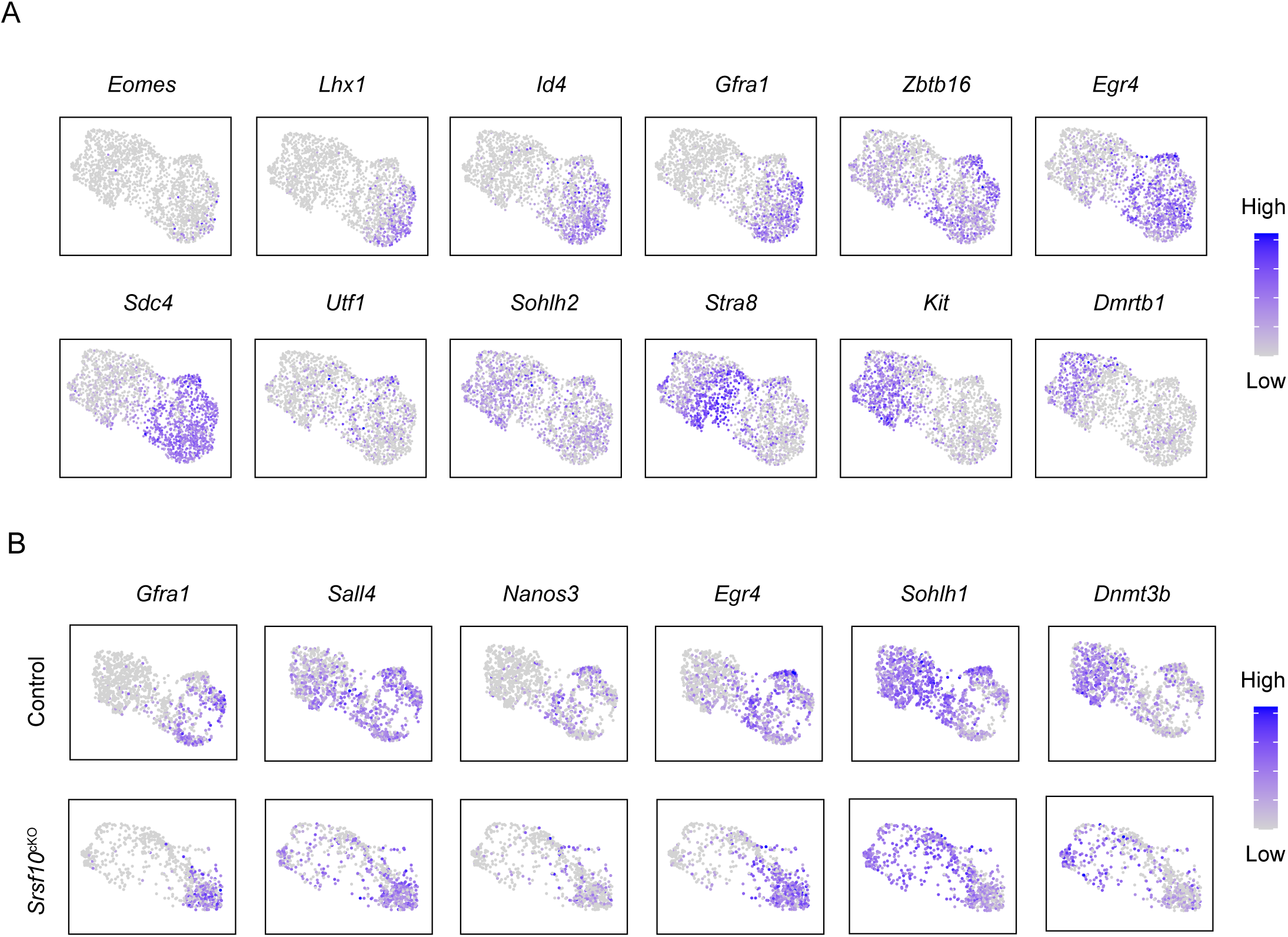
The expression pattern of spermatogonial maker genes in the subtypes of control and *Srsf10*^cKO^ samples. A. Gene expression patterns of marker genes corresponding to each cellular state on the UMAP plots. B. UMAP plots of the expression patterns of selected marker genes in each subtype in control and *Srsf10*^cKO^ samples.

**Figure 6-figure supplement 1.**
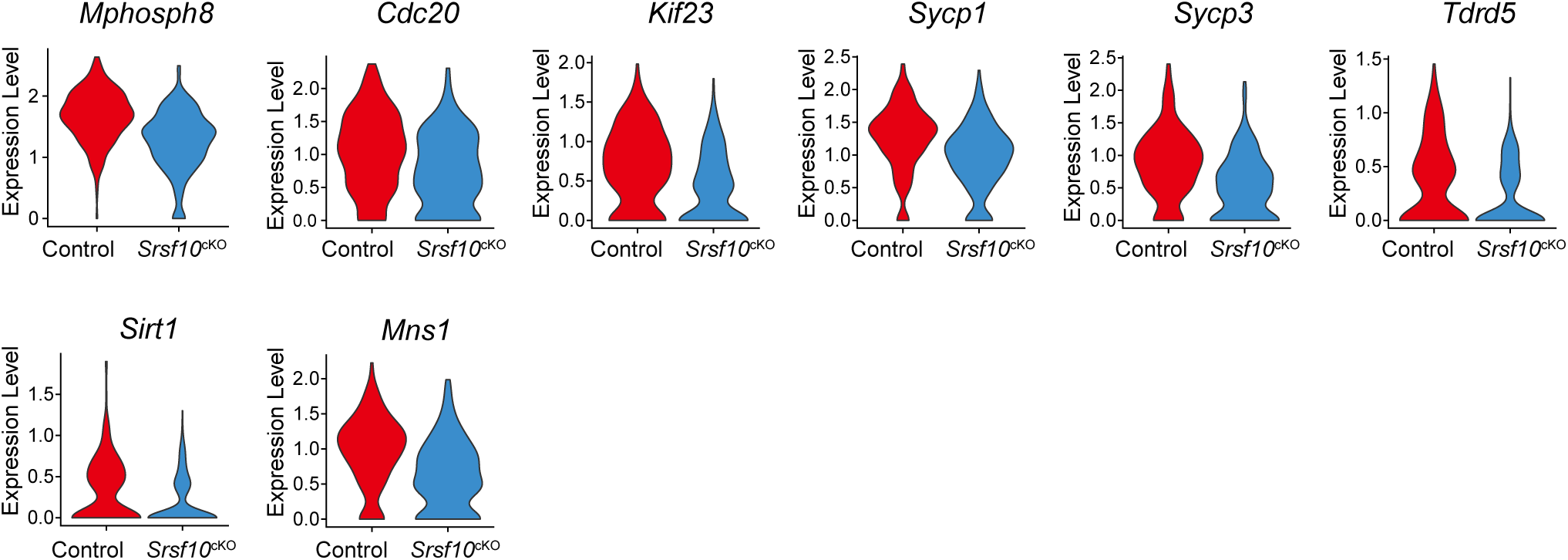
The expression pattern of key genes of the USSC1 subtype in control and *Srsf10*^cKO^ groups. The violin plots show the expression level of cell cycle key genes and spermatogenesis-related genes of the USSC1 subtype in control and *Srsf10*^cKO^ groups.

**Figure 6-figure supplement 2.**
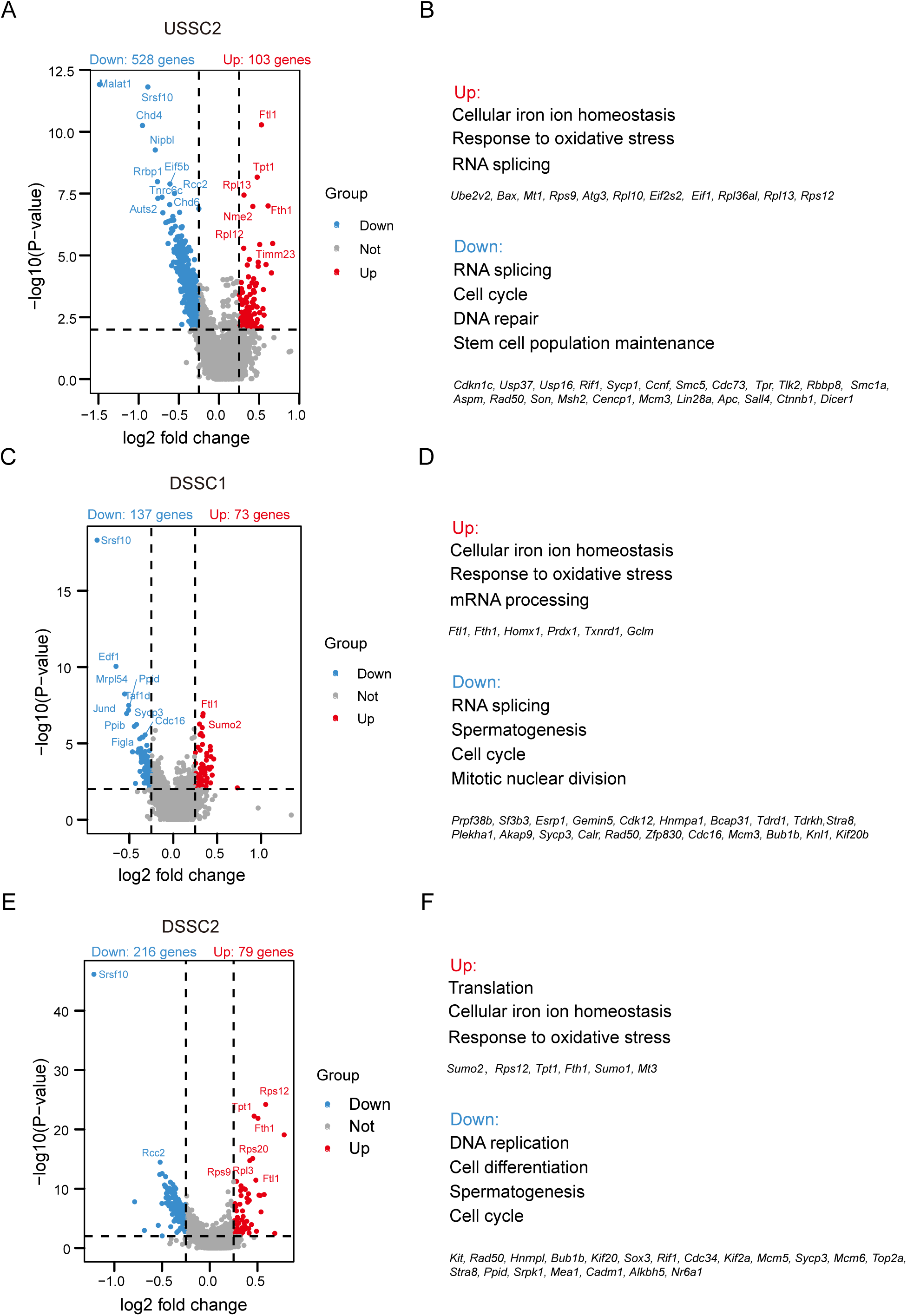
Identification of the differential expression genes in USSC2, DSSC1 and DSSC2 subtypes between control and *Srsf10*^cKO^ samples. A, C and E. Volcano plot of significantly differentially expressed transcripts of the USSC2, DSSC1 and DSSC2 subtype, respectively, in *Srsf10*^cKO^ sample compared with the control. Blue dots represent significantly down-regulated transcripts, while red dots show significantly up-regulated transcripts (Log2 fold change >= 0.25, < 0.01). Grey dots illustrated unchanged transcripts. B, D and F. Gene ontology of up-regulated and down-regulated genes in *Srsf10*^cKO^ USSC2, DSSC1 and DSSC2 subtype, respectively.

**Figure 6-source data 1** Quantification of the ratio of KI67^+^PLZF^+^ cells, EdU^+^PLZF^+^ cells and CAP3^+^PLZF^+^ cells in PLZF^+^ cells in control and *Srsf10*^cKO^ testes.

**Figure 7-figure supplement 1.**
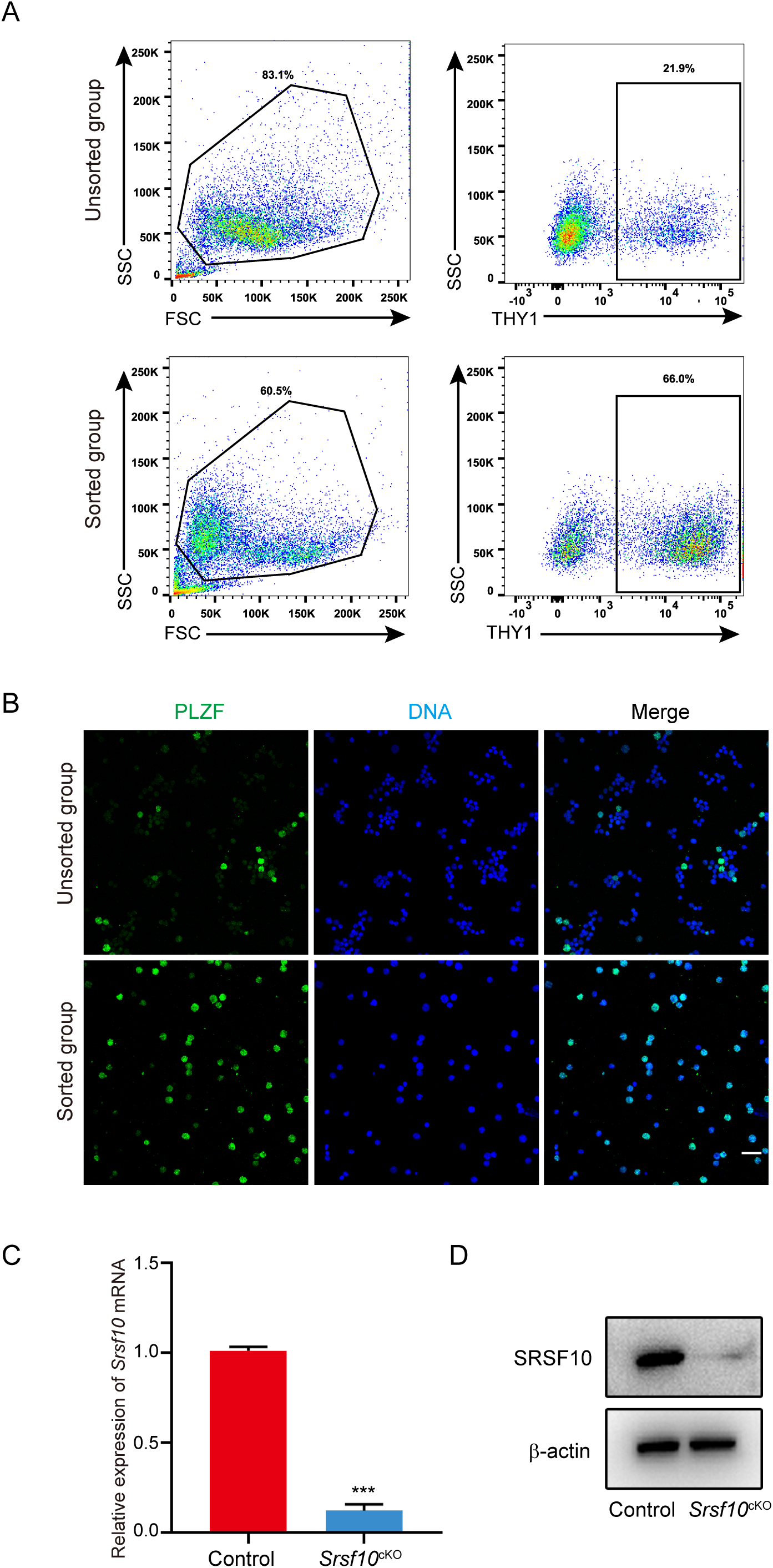
Sorting strategies for THY1^+^ spermatogonia in control and *Srsf10*^cKO^ testes at P6. A. Gating strategy for identifying THY1^+^ spermatogonia from P6 testes. FSC vs SSC gating was used to clean up the debris and dead cells. The percentage of THY1^+^ cells in the unsorted and sorted groups corresponds to 21.9 and 66.0%, respectively. B. Immunofluorescence staining for PLZF in the unsorted and sorted groups. Scale bar, 20 µm. C. Quantitative RT-PCR analysis of *Srsf10* expression in the sorted THY1^+^ spermatogonia from control and *Srsf10*^cKO^ testes at P6. -actin was used as the internal control. *** < 0.001, Error bars represent s.e.m. D. Western blot analyses of SRSF10 expression in the sorted THY1^+^ spermatogonia from control and *Srsf10*^cKO^ testes at P6. -actin was used as the loading control.

**Figure 7-figure supplement 2.**
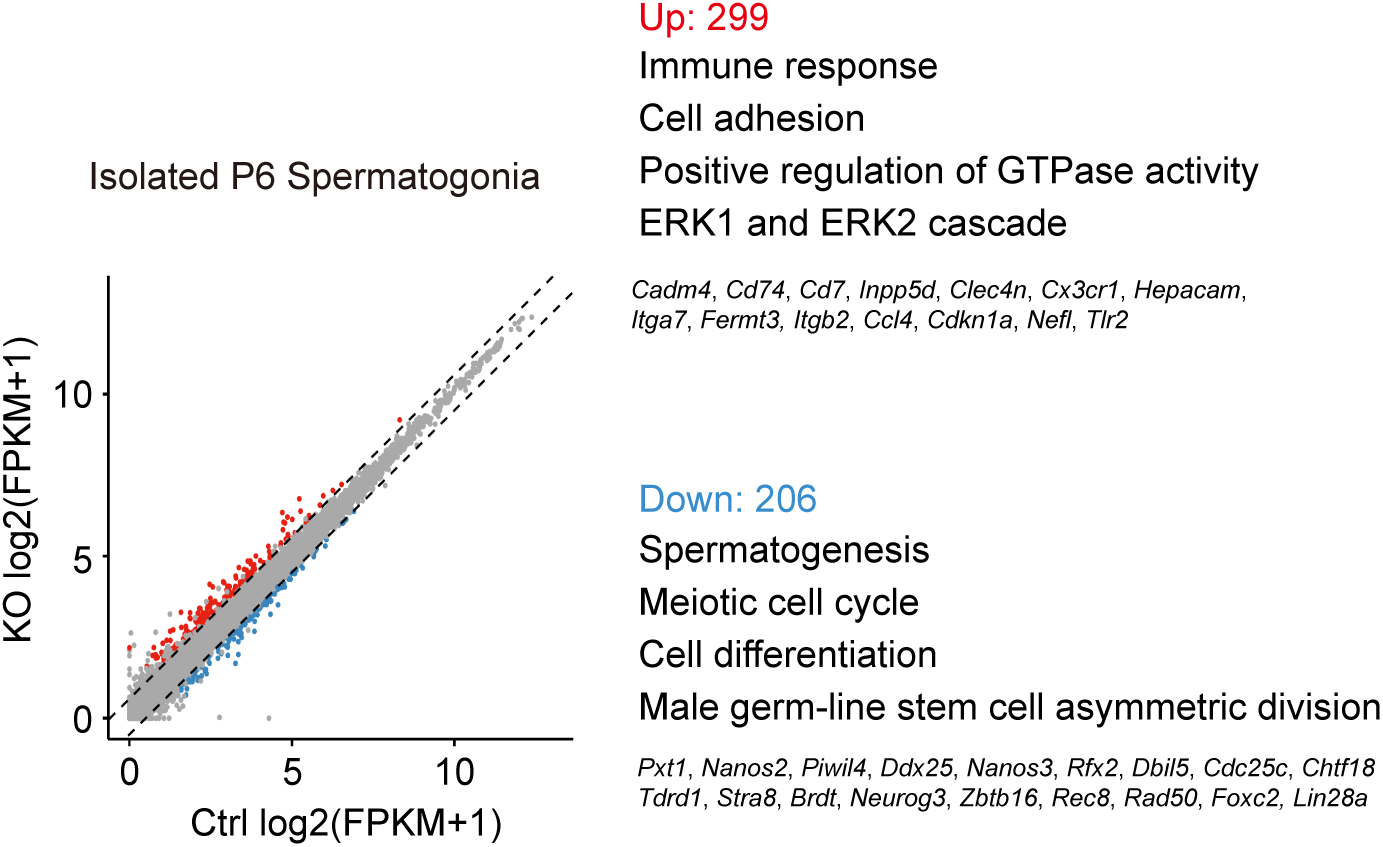
Analysis of the transcriptome of THY1^+^ spermatogonia enriched from control and *Srsf10*^cKO^ testes at P6. A. Scatter plots showing the expression of genes in THY1^+^ spermatogonia enriched from control and *Srsf10*^cKO^ testes at P6. Blue dots represent significantly down-regulated genes, while red dots show significantly up-regulated genes (FPKM >= 2, fold change >= 1.5, < 0.01). Grey dots represent unchanged genes. B. Gene ontology of up-regulated and down-regulated genes in *Srsf10*^cKO^ THY1^+^ spermatogonia.

**Figure 7-figure supplement 3.**
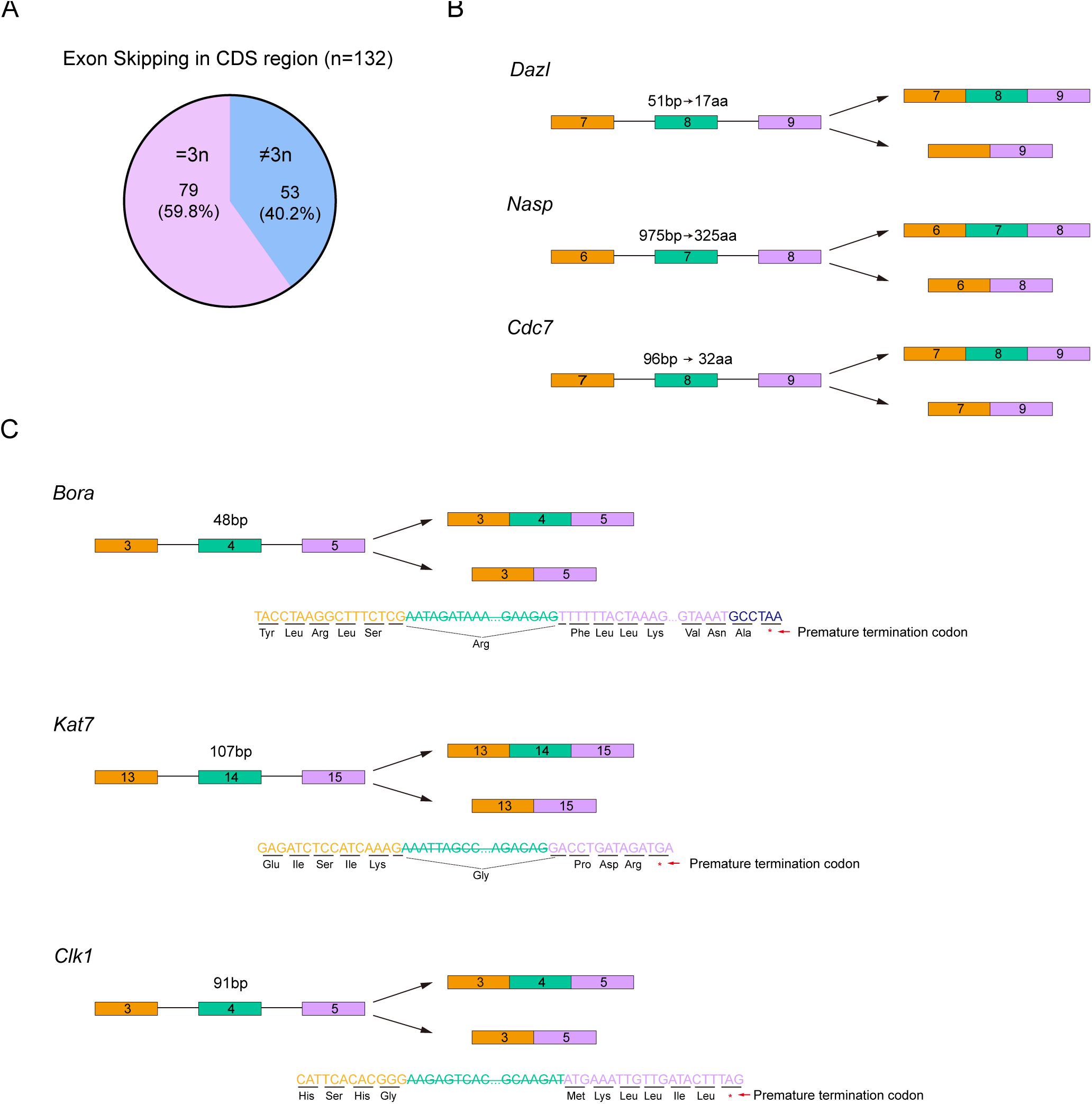
The possible results of abnormal exon skipping in *Srsf10* depleted spermatogonia. A. Statistically analysis of the base number of the excluded or included exons (multiple of 3, =3n and not multiple of 3, ≠3n). B. Take *Dazl*, *Nasp* and *Cdc7* for example to show the outcomes of abnormal skipped 3n base number exons. C. Take *Bora*, *Kat7* and *Clk1* for example to analyze the outcomes of abnormal skipped ≠3n base number exons.

**Figure 7-figure supplement 4.**
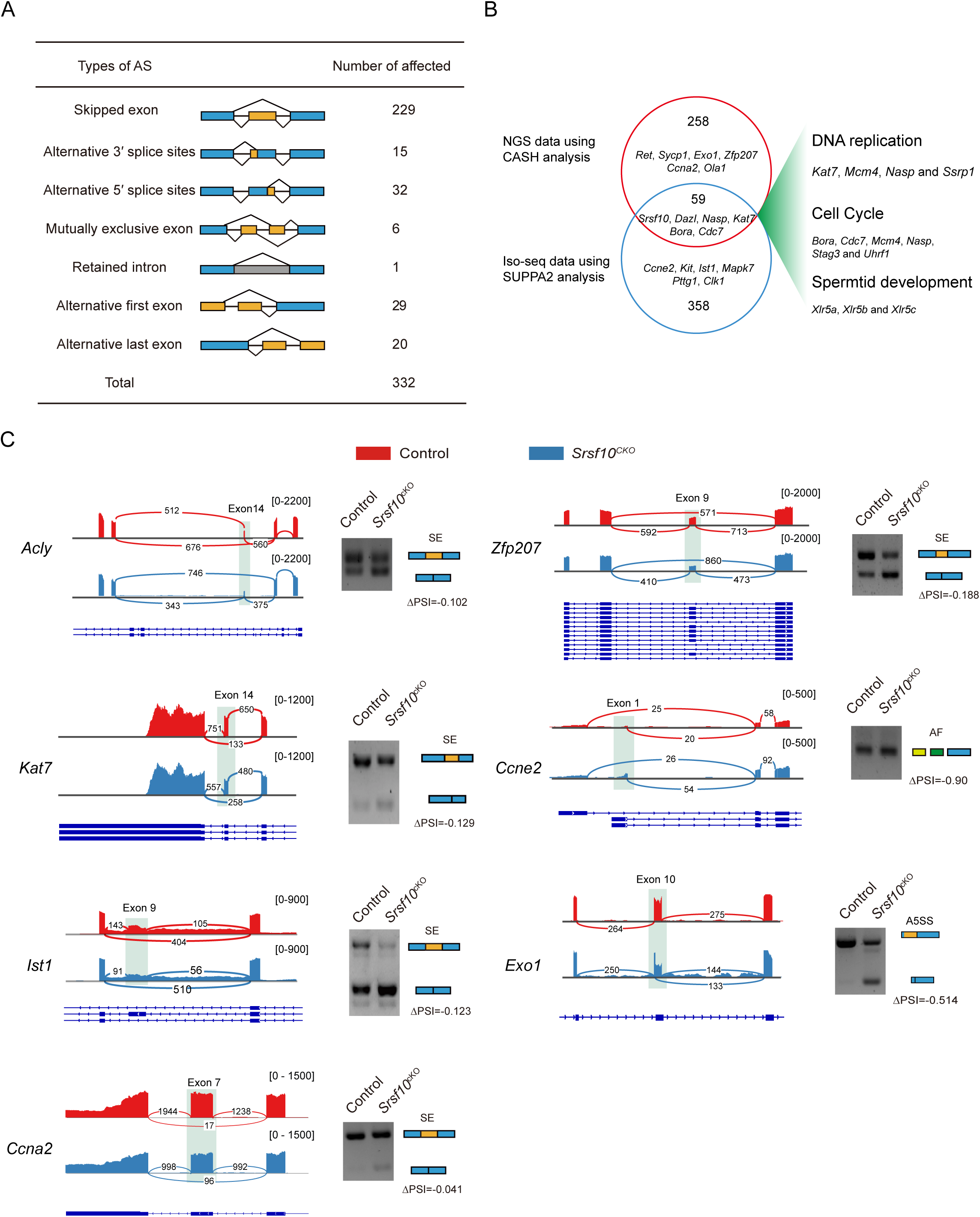
Visualization and validation of some differentially spliced functional genes in control and *Srsf10* depleted spermatogonia. A. Seven AS events were significantly affected by depletion of *Srsf10* in the NGS data using CASH software analysis. ( <0.05). B. Venn diagram depicting the overlap of significantly affected AS genes between NGS data using CASH and Iso-seq data using SUPPA2. Gene ontology (GO) terms of the 59 shared genes involved in specific functional terms which were shown on the right. C. Tracks from IGV are shown for selected candidate genes (left). Differentially spliced exons are shaded. Schematics of alternative splicing events are shown (blue and yellow rectangles) (right). Changes in “percent spliced in (PSI)” between control and *Srsf10*^cKO^ spermatogonia are shown below splicing schematics (ΔPSI). SE, Skipped exon, A5SS, Alternative 5’ splice sites, A3SS, Alternative 3’ splice sites, AF, Alternative first exon, AL, Alternative last exon.

**Figure 7-source data 1** Differential alternative splicing events in the Iso-seq data using SUPPA2 analysis.

**Figure 7-source data 2** Relative expression of *Kit* in control and *srsf10*^cKO^ spermatogonia.

**Figure 7-figure supplement 1-source data 1** Relative mRNA expression of *Srsf10* in enriched cells.

**Figure 7-figure supplement 3-source data 1** Analysis the skipped exon events.

**Figure 7-figure supplement 4-source data 1** Differential alternative splicing events in the NGS-seq data using CASH analysis.

**Table 1-source data 1:** The fertility of Srsf10cKO male mice.

